# Functional Activity of HIV-1 bNAbs Across Diverse Strains is Driven by Binding Site and Can be Enhanced Through Fc Engineering

**DOI:** 10.1101/2025.11.12.688078

**Authors:** Chia Jung Li, Eunice Lim, Meredith Phelps, Jacqueline M. Brady, Vintus Okonkwo, Caitlin McCarthy, Caroline Alexander, Margaret E. Ackerman, Daniel Lingwood, Alejandro B. Balazs

## Abstract

Broadly neutralizing antibodies (bNAbs) are promising tools for HIV-1 treatment and prevention, due to their ability to mediate both Fab-dependent neutralization and Fc-dependent effector functions. While antibody-dependent cellular cytotoxicity (ADCC) and antibody-dependent cellular phagocytosis (ADCP) have been implicated in antiviral activity, the extent to which these functions vary across bNAb epitopes, viral strains, and Fc region remains unclear. Here, we systematically evaluated twenty bNAbs targeting five distinct Env epitopes against a diverse panel of nine HIV-1 strains. We found that epitope specificity and Env sequence both critically influenced bNAb effector functions. CD4 binding site (CD4bs)- and V3 glycan-targeting bNAbs mediated the broadest and most potent ADCC and ADCP, while V1/V2 apex-directed bNAbs preferentially induced ADCP. In contrast, MPER-targeting bNAbs triggered ADCC more selectively, and gp120/gp41 interface-targeting bNAbs showed limited activity. Irrespective of binding epitope, different strains exhibited a broad range of effector function sensitivities. To investigate the impact of Fc modifications on this variability, we subclass-switched and introduced previously identified Fc mutations known to change Fcγ receptor affinity. Some of these mutations significantly boosted ADCC, while IgG3 subclass switching dramatically enhanced ADCP, even against highly resistant strains. Collectively, these results demonstrate that effector function is shaped by both antibody specificity and viral Env context and that rational Fc modification has the potential to improve antibody-based therapeutics against HIV.

**One Sentence Summary:** This study systematically evaluates the impact of bNAb epitope specificity, Env sequence, and Fc engineering on antibody-mediated effector functions.

## INTRODUCTION

Human immunodeficiency virus type 1 (HIV-1) remains a global health burden, with over 39 million people currently living with the virus and approximately 1.3 million new infections reported each year (UNAIDS, 2023). Although antiretroviral therapy (ART) effectively suppresses viral replication and halts disease progression, it does not eliminate the virus (*1*). A key barrier to a cure is the persistence of long-lived reservoirs of latently infected cells that harbor integrated proviral DNA and evade immune surveillance. These reservoirs have an estimated half-life of ∼44 months and are capable of reinitiating systemic viremia if ART is interrupted (*2*). Consequently, lifelong treatment is required, and alternative strategies that can eliminate or durably control HIV in the absence of ART are urgently needed. Eliminating infected cells and latent reservoirs is essential to achieving a functional cure. However, while ART blocks new infection, it does not activate immune mechanisms to clear cells already harboring viruses. Broadly neutralizing antibodies (bNAbs) have emerged as promising therapeutic candidates due to their ability to recognize conserved epitopes on the HIV-1 envelope glycoprotein (Env) and neutralize a wide range of viral strains (*3–5*). These bNAbs target key Env regions, including the V1/V2 apex, V3 glycan supersite (V3 glycan), CD4 binding site (CD4bs), gp120/gp41 interface, and membrane-proximal external region (MPER) (*6*). Roughly 10–25% of individuals with HIV naturally develop bNAbs over time, though fewer than 1% develop “elite” bNAbs capable of cross-clade neutralization (*7*).

Beyond neutralization, many bNAbs mediate Fc-dependent effector functions that contribute to antiviral activity *in vivo*. These include antibody-dependent cellular cytotoxicity (ADCC), antibody-dependent cellular phagocytosis (ADCP), and complement-dependent cytotoxicity (CDC), all of which enable immune cells to recognize and eliminate virus or infected cells (*8*). In ADCC, antibodies bind to HIV-infected cells and engage Fc gamma receptor 3a (FcγRIIIa, also known as CD16) on natural killer (NK) cells, triggering cytotoxic granule release and target cell lysis (*9*). In ADCP, antibody-opsonized targets are engulfed by phagocytes via FcγR-mediated uptake (*10*). These mechanisms have been associated with protective immunity and viral control (*11–13*), and may contribute to vaccine-induced protection (*14, 15*). For instance, ADCC correlated with delayed disease progression and partial protection in the RV144 vaccine trial (*11, 16*), and was inversely associated with infection risk in the Vax004 vaccine trial (*17*). Similarly, IgG3 antibodies, which possess extended hinge regions, can more effectively engage Fcγ receptors and stimulate cellular activity(*18*). Consistent with this, IgG3 responses have been associated with enhanced antiviral activity and reduced risk of infection in vaccine studies (*14, 19, 20*). In our prior study using a humanized mouse model, VRC07 lost protective efficacy when its Fc domain was switched from IgG1 to IgG2, which reduced ADCC and ADCP activity (*21*). Another study also showed that b12 (a CD4bs-targeting bNAb) which was engineered to lack innate functions failed to protect non-human primates despite intact neutralization (*22*). However, introducing similar mutations into the more potent bNAb PGT121 did not impair protection (*23*). These differing results underscore the complexity of Fc-mediated protection and raise questions about the factors that govern effector function (*24*).

Epitope specificity has been proposed as a key variable influencing Fc function (*25*). Factors such as epitope exposure, antibody orientation, and Fc accessibility may also impact ADCC and ADCP efficiency (*26–28*). However, findings across studies have been inconsistent. For example, some studies report that infected primary cells are more effectively eliminated by V1/V2 apex-targeting bNAbs such as PG9 and PG16, but not by CD4bs antibodies (*25*)—a pattern that contrasts with findings in cell line-based models (29). Other reports show that CD4bs bNAbs, such as 3BNC117, can mediate robust ADCC against reactivated primary cells from ART-treated individuals (*30*). These discrepancies may reflect differences between Env proteins or in experimental model systems, emphasizing the need for systematic analyses across multiple epitopes and viral strains.

The Fc domain is known to be a critical determinant of effector activity for antibodies (*31*). Prior work examining HIV bNAbs of different subclasses found that IgG1 and IgG3 are strong inducers of Fc-mediated effector mechanisms, with IgG3 being particularly effective in mediating ADCP (*32, 33*), while IgG2 and IgG4 generally induce more subtle responses (*34*). HIV bNAb Fc differences, including point mutations (*35*), subclass switching (*32*), and glycosylation (*36*), can impact FcγR binding and downstream activity. For example, afucosylation increases FcγRIIIa affinity and enhances ADCC (*27, 37*), GASDALIE mutations improve NK cell activation (*38*), and IgG3 subclass switching or hinge elongation enhances ADCP (*39–41*) and *in vivo* activity (*42*). Despite these advances, how such Fc modifications influence bNAb function across diverse HIV-1 Envs and antibodies remains poorly defined.

To address these questions, we conducted a systematic analysis of bNAb-mediated effector functions across twenty bNAbs targeting five distinct Env epitopes on cell lines expressing a panel of primary HIV-1 Envs that represent global diversity (*43*). We assessed neutralization, Env binding, ADCC, and ADCP for each bNAb–virus pair to examine how epitope specificity and Env sequence shape antibody function. Additionally, we engineered the CD4bs-directed bNAb N49P7 and V3 glycan-targeting bNAb PGT121 with Fc domains previously shown to enhance or decrease FcγR engagement, and tested their impact on effector activity and neutralization potency. Our results revealed epitope-and Env sequence-dependent variation in bNAb function and demonstrated that Fc engineering can selectively enhance ADCC and ADCP, offering insights for the rational design of antibody-based therapies to promote immune-mediated clearance of HIV-infected cells.

## RESULTS

### Effector functions of bNAbs were determined by targeted viral epitopes

To investigate the influence of epitope specificity on effector function, we selected a panel of twenty HIV-1 bNAbs targeting five major Env vulnerability sites (**Table S1**). Each bNAb was expressed as an IgG1 and classified by epitope specificity: V1/V2 apex, V3 glycan, CD4bs, gp120/gp41 interface, or MPER. This bNAb panel was first assessed for neutralization potency against replication-competent HIV_REJO.c_, a clade B transmitted/founder (T/F) virus, revealing that bNAbs targeting the V1/V2 apex and CD4bs epitopes exhibited the highest neutralization potency, whereas V3 glycan- and MPER-targeting bNAbs displayed lowest potency (**Figure 1A, Figure S1**).

**Figure 1.**
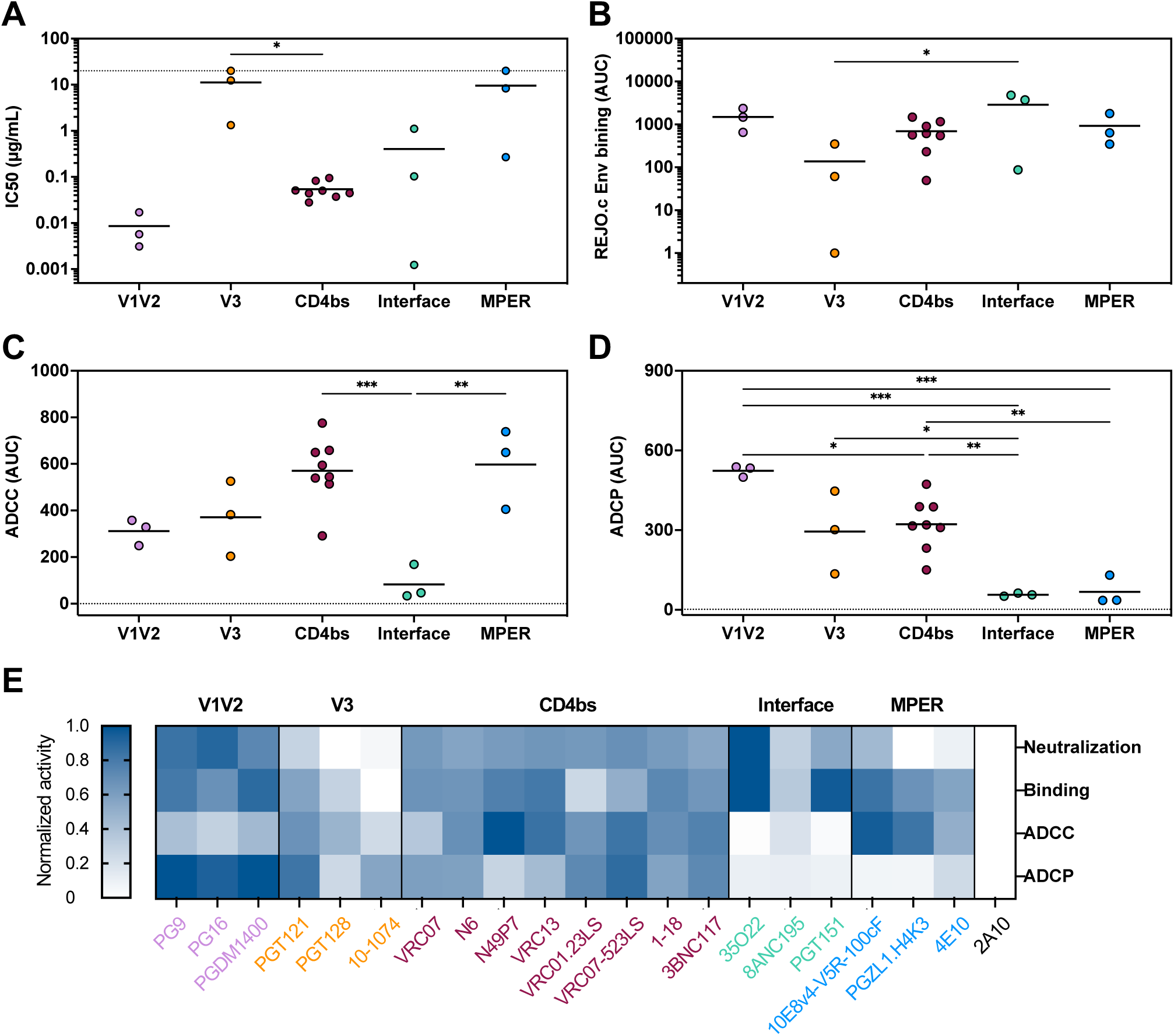
Effector functions of bNAbs vary by HIV_REJO.c_ Env epitope. A panel of twenty HIV broadly neutralizing antibodies (bNAbs), targeting five distinct epitopes, was selected for functional analysis against HIV_REJO.c_. (**A**) Neutralization potency of the bNAb panel, measured by IC_50_ against replication-competent HIV_REJO.c_. Data were pooled by epitope, with each dot representing an individual bNAb. (**B-D**) Functional characterization of the bNAb panel against HIV_REJO.c_ Env-expressing CEM.NKr cells are colored by epitope specificity. Env binding (B), ADCC (C), and ADCP (D) are represented by scatter plots. Horizontal bars represent the mean value for each epitope. Statistical significance was determined using one-way ANOVA followed by Tukey’s multiple comparisons test (*p < 0.0322, **p < 0.0021, ***p < 0.0002, ****p < 0.0001). (**E**) Heatmap depicting the functional profiles of bNAbs against HIV_REJO.c_. Data is presented as -log(IC_50_) for neutralization, log-transformed AUC for binding, and AUC for ADCC and ADCP, normalized within each function. A value of 1 represents the highest activity, and 0 represents the lowest.

To create a uniform target cell line for functional assays, we used a lentiviral vector to stably express codon-optimized Env from HIV_REJO.c_ in CEM.NKr cells, an immortalized T cell line. The construct included ZsGreen as a fluorescent marker to facilitate target cell identification (**Figure S2A**). To confirm surface expression of Env, we constructed a plasmid encoding the same codon-optimized Env fused to a C-terminal Neon fluorescent tag. Fluorescence microscopy of HEK293T cells transfected with this construct confirmed surface localization of Env (**Figure S2B, left**), whereas cells transfected with Neon alone exhibited diffuse cytoplasmic fluorescence (**Figure S2B, right**).

Next, we assessed bNAb binding to Env-expressing CEM.NKr cells and observed uniform but distinct binding profiles for each antibody (**Figure S3A**). To compare relative binding, we measured bound antibody across a range of antibody concentrations of each bNAb and found that interface-targeting bNAbs exhibited the highest binding area under the curve, while V3 glycan-targeting bNAbs showed limited binding (**Figure 1B, Figure S3B**). These Env-expressing cells were then used to evaluate ADCC and ADCP. For ADCC, we generated CD16.NK-92 cells—a natural killer (NK) cell line engineered to express FcγRIIIa (also known as CD16), a receptor that binds the Fc region of antibodies and initiates downstream effector functions—to facilitate NK cell-mediated killing of target cells (**Figure S4A**). ZsGreen+ Env-expressing CEM.NKr and ZsGreen-parental CEM.NKr cells were mixed at a 1:1 ratio, incubated with bNAbs, and co-cultured with CD16.NK-92 cells. Target cell lysis was quantified by flow cytometry after 6 hours (**Figure S4B, C**). The data demonstrates that CD4bs- and MPER-targeting bNAbs mediated the strongest ADCC responses, while interface-directed bNAbs were largely inactive. V1/V2 apex-directed bNAbs showed moderate potency, and V3 glycan-targeting bNAbs exhibited variable activity (**Figure 1C, Figure S4D**). For ADCP, THP-1 monocytes were used as effector cells to measure phagocytosis of Env-expressing targets (**Figure S5A, B**). V1/V2 apex-targeting bNAbs induced the most robust ADCP, whereas interface- and MPER-targeting bNAbs showed limited activity (**Figure 1D, Figure S5C**).

To compare the activity of functions of each bNAb, neutralization, binding, ADCC, and ADCP activities were normalized within each function and shown as a heatmap (**Figure 1E**). CD4bs-targeting bNAbs potently triggered both ADCC and ADCP, while V1/V2 apex-directed bNAbs primarily mediated ADCP. MPER-directed bNAbs were more effective at ADCC than ADCP, and interface-targeting bNAbs were largely ineffective in both assays despite strong Env binding.

### Epitope specificity dictated bNAb-mediated effector functions across diverse strains

Following our initial evaluation of HIV_REJO.c_, we expanded the analysis to nine HIV-1 strains spanning clades A, AC, B, CRF01_AE, CRF07_BC, C, and G (**Table S2**). These strains were selected to represent the global diversity of circulating strains (*43*). To explore viral sequence similarity and clade classification, we constructed a phylogenetic tree and calculated pairwise amino acid identity among the Env sequences (**Figure 2A, B**). To assess bNAb-mediated neutralization across these strains, chimeric HIV constructs were generated by replacing the Env of HIV_JR-CSF_ with the Env sequences of HIV_398F1_, HIV_246F3_, HIV_CNE55_, HIV_CH119_, HIV_25710_, HIV_CE1176_, or HIV_X1632_. We then measured neutralization potency (IC_50_) of the bNAb panel against HIV_REJO.c_, HIV_JR-CSF_, and each of the chimeras. In this assay, bNAbs targeting the V1/V2 apex and CD4bs epitopes showed strong neutralization, while those targeting the gp120/gp41 interface or MPER epitopes showed weaker potency (**Figure 2C, Figure S6**).

**Figure 2.**
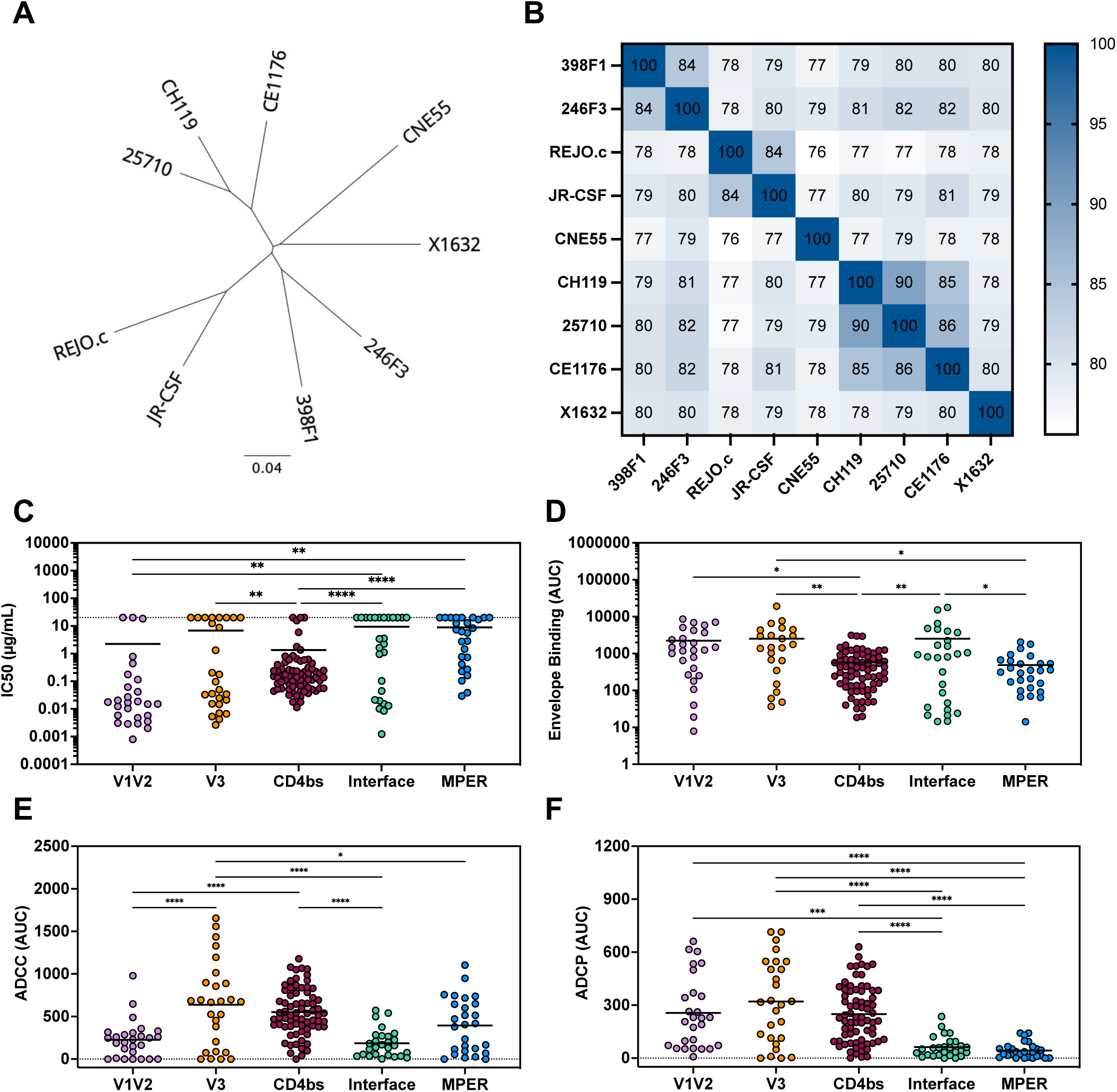
Effector functions across diverse HIV strains are dependent on Env epitope. (**A**) Phylogenetic tree of HIV-1 strains based on Env amino acid sequences, constructed using the neighbor-joining method. The scale bar indicates 0.04 amino acid substitutions per site. (**B**) Pairwise percentage amino acid identity matrix of HIV Env sequences. (**C**) Neutralization potency (IC_50_) of bNAbs against diverse viruses. Scatter plots display pooled data from multiple bNAbs targeting the same epitope. (**D-F**) Functional activity of the bNAb panel against CEM.NKr cells expressing Env from nine HIV-1 strains are colored by epitope specificity. Env binding (D), ADCC (E), and ADCP (F) are represented by scatter plots. Horizontal bars indicate mean values. Statistical significance was determined by one-way ANOVA followed by Tukey’s multiple comparisons test (*p < 0.0322, **p < 0.0021, ***p < 0.0002, ****p < 0.0001).

Following the same approach used for HIV_REJO.c_, we codon-optimized each Env and designed constructs with a C-terminal Neon fusion. Fluorescence imaging of transfected HEK293T cells confirmed surface localization of Env proteins (**Figure S7A**). Quantification of Neon fluorescence revealed that Env proteins from all HIV-1 strains exhibited comparable expression levels (**Figure S7B**). Each Env was then stably expressed on CEM.NKr cells using the same expression plasmid design as for HIV_REJO.c_. Antibody binding and functionality were measured using the same flow-based assays. Flow cytometry analysis revealed that bNAbs targeting the V1/V2 apex, V3 glycan, and gp120/gp41 interface exhibited high levels of Env binding, whereas CD4bs- and MPER-targeting bNAbs displayed poorer binding efficiency (**Figure 2D, Figure S8**). Effector function assays demonstrated distinct patterns of bNAb activity. In ADCC assays, V3 glycan-, CD4bs, and MPER-targeting bNAbs facilitated potent target cell elimination by NK cells, whereas V1/V2 apex- and gp120/gp41 interface-directed bNAbs exhibited low ADCC activity (**Figure 2E, Figure S9**). In ADCP assays, bNAbs targeting the V1/V2 apex, V3 glycan, and CD4bs epitopes exhibited strong ADCP activity, whereas gp120/gp41 interface- and MPER-targeting bNAbs demonstrated weak ADCP activity (**Figure 2F, Figure S10**). Collectively, V3 glycan- and CD4bs-targeting bNAbs mediated both ADCC and ADCP across diverse Envs. V1/V2 apex-directed bNAbs preferentially triggered ADCP, while MPER-targeting bNAbs mainly induced ADCC. Interface-targeting bNAbs exhibited minimal effector functions across all tested strains. These findings emphasize the role of epitope specificity in shaping bNAb function across genetically distinct Envs.

### Different strains exhibited variable bNAb-mediated effector functions

To visualize individual antibody performance across Envs, we normalized the neutralization, binding, ADCC, and ADCP data and displayed them as a heatmap (**Figure 3**). This analysis reveals substantial functional variability across different strains, even among bNAbs targeting the same epitope. For example, although PG9 and PG16 both recognize the V1/V2 apex and share similar neutralization profiles, they differed in effector activity—PG9 consistently mediated robust ADCP across all strains, whereas PG16 showed more limited activity. Among V3 glycan-targeting bNAbs, binding levels generally correlated with both ADCC and ADCP activity, suggesting that strong Env engagement was a key determinant of effector function. However, notable exceptions were observed; for example, 10-1074 triggered ADCC and/or ADCP despite weak binding to HIV_246F3_ and HIV_REJO.c_ Envs, while PGT121 demonstrates broad Env binding but inconsistent effector functions, underperforming against cells expressing HIV_246F3_ Env, HIV_CNE55_ Env, and HIV_X1632_ Env. Interestingly, PGT128 failed to neutralize HIV_REJO.c_ and HIV_CNE55_ yet retained the capacity to mediate both ADCC and ADCP. Compared to the strain-dependent but potent effector activity of V3 glycan-targeting bNAbs, CD4bs-targeting bNAbs consistently mediated both ADCC and ADCP across all tested strains. N49P7, however, exhibited weaker ADCP despite moderate Env binding. In contrast, gp120/gp41 interface-targeting bNAbs such as 35O22 and PGT151 showed strong Env binding but failed to induce either function, while ADCC activity of MPER-targeting bNAbs was strain-dependent.

**Figure 3.**
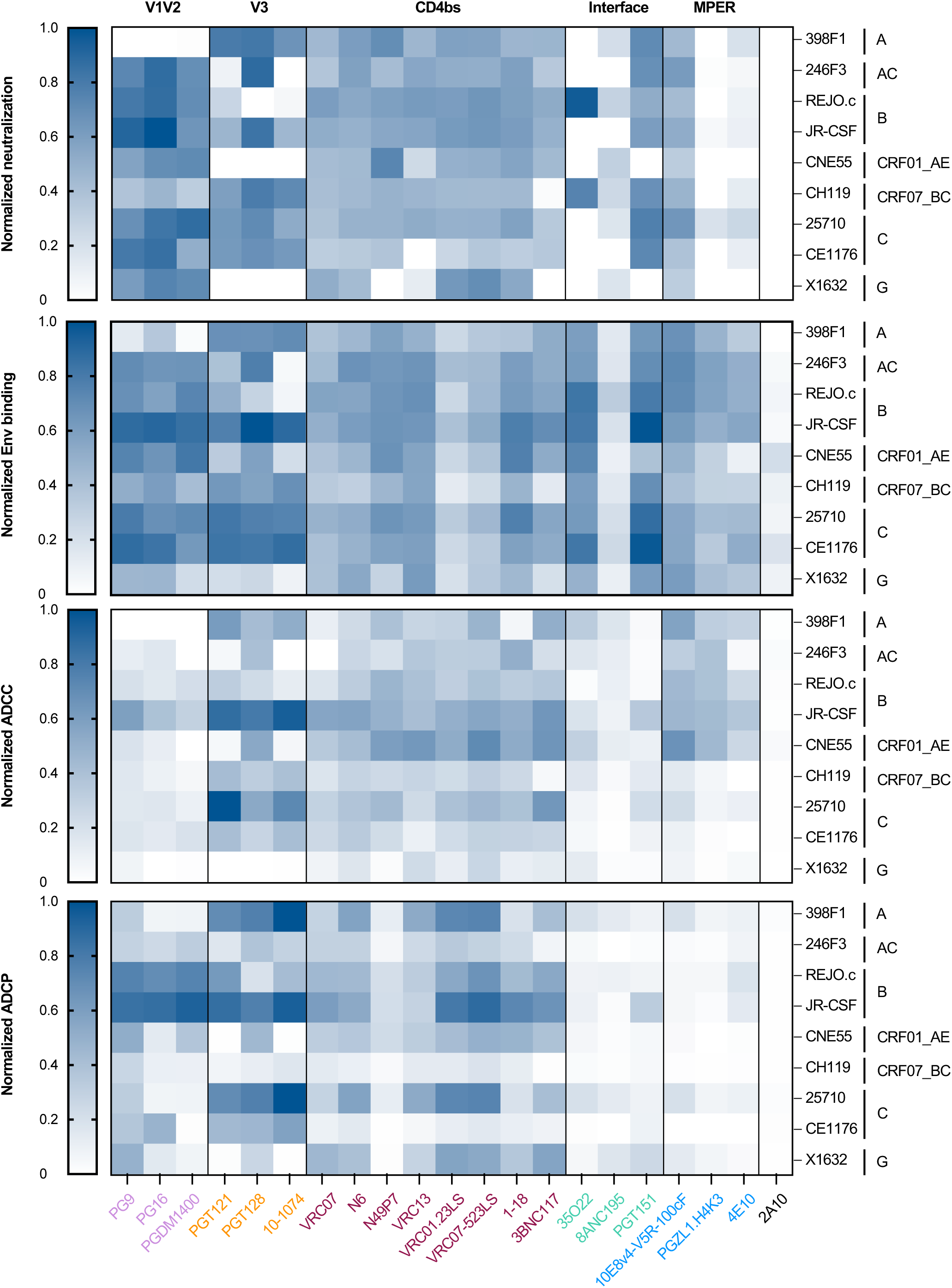
Antibody-mediated effector functions vary by strain. Heatmap showing the functional profiles of bNAbs against the HIV-1 panel. Data are displayed as -log10(IC_50_) for neutralization, log10-transformed AUC for Env binding, and AUC for ADCC and ADCP. All values were normalized on a 0 to 1 scale within each function. bNAbs were grouped and color-coded by epitope specificity. HIV clades are indicated on the heatmap. Viral Envs included HIV_398F1_ (clade A), HIV_246F3_ (clade AC), HIV_REJO.c_ (clade B), HIV_JR-CSF_ (clade B), HIV_CNE55_ (clade CRF01_AE), HIV_CH119_ (clade CRF07_BC), HIV_25710_ (clade C), HIV_CE1176_ (clade C), and HIV_X1632_ (clade G).

We also observed Env-dependent differences in sensitivity to effector responses. Cells expressing HIV_X1632_ Env (clade G) were broadly resistant to ADCC, whereas HIV_CH119_ Env (clade CRF07_BC) showed reduced susceptibility to ADCP. In contrast, HIV_JR-CSF_ Env (clade B) and HIV_25710_ Env (clade C) were highly sensitive to both ADCC and ADCP (**Figure 3**). Env sequence identity alone did not consistently predict effector function sensitivity (**Figure 2B**); for instance, HIV_25710_ and HIV_CE1176_ Envs (both clade C) were closely related but differed in ADCP susceptibility, and HIV_REJO.c_ and HIV_JR-CSF_ Envs (both clade B) showed similar sequence identity yet exhibited distinct ADCC responses. These observations suggest that sequence similarity or clade assignment is not a reliable predictor of effector function activity. Instead, Env structural features and epitope accessibility likely play a more dominant role in shaping susceptibility to bNAb-mediated effector functions.

To better understand the relationship between bNAb functions within one strain, we generated heatmaps for each individual HIV-1 Env (**Figure S11**). The data revealed that all V1/V2 apex-targeting bNAbs exhibited weak binding to HIV_398F1_ Env and failed to neutralize this strain; however, PG9 remained capable of mediating ADCP. Notably, strong neutralization and effector activities were observed for V3 glycan-targeting bNAbs against HIV_398F1_, HIV_JR-CSF_, HIV_25710_, and HIV_CE1176_, highlighting their therapeutic potential against these strains. While CD4bs-targeting bNAbs broadly mediated both ADCC and ADCP across all tested strains, the magnitude of ADCC was higher against HIV_CNE55_, whereas ADCP was higher against HIV_398F1_, HIV_JR-CSF_, HIV_25710_, and HIV_X1632_. Together, these findings underscore the variability of antibody effector responses across diverse HIV-1 Envs and the influence of both epitope specificity and Env context on bNAb function.

### Fc-mediated effector functions correlated with Env binding and neutralization potency

To examine the relationships between bNAb-mediated functions and Env binding, we performed Spearman correlation analyses. Significant positive correlations were observed between Env binding and ADCC (r = 0.247, P = 0.0008) or ADCP (r = 0.271, P = 0.0002), suggesting that stronger Env engagement generally enhanced Fc-mediated effector responses. However, there were some V3 glycan-targeting bNAbs-Env combinations that showed relatively poor binding but potent effector functions (**Figure 4A, B**). As expected, Env binding also showed a significant positive correlation with neutralization potency (r = −0.198, P = 0.0076) (**Figure 4C**). Finally, ADCC and ADCP exhibited a strong positive correlation (r = 0.557, P < 0.0001), indicating that bNAbs capable of inducing one effector function were likely to trigger the other, potentially reflecting shared mechanisms or similar FcγR engagement (**Figure 4D**).

**Figure 4.**
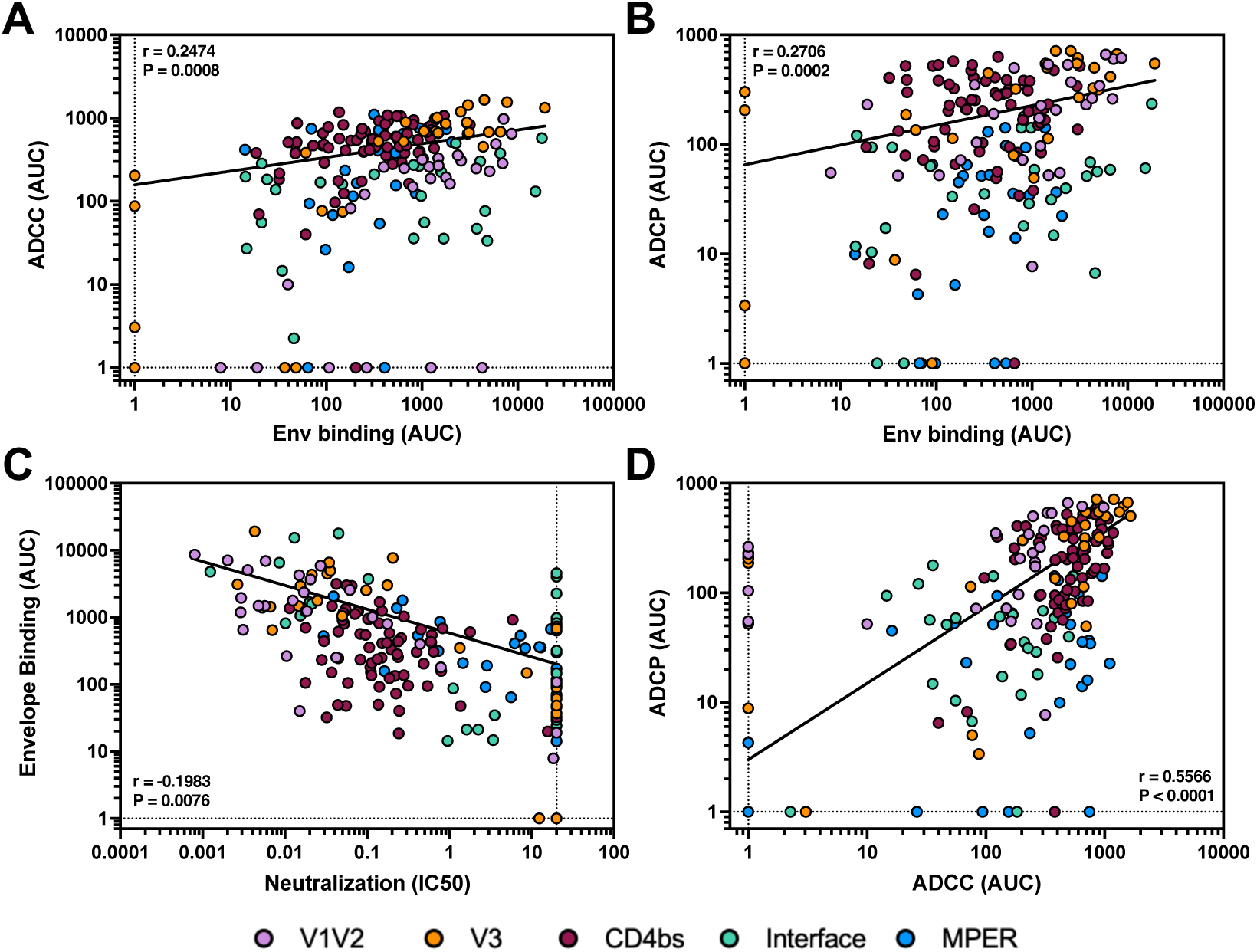
Functional activity correlates with Env binding. (**A** and **B**) Scatter plot correlations between bNAb-mediated ADCC (A) or ADCP (B) and Env binding were evaluated across multiple HIV-1 strains. Each data point represents a single bNAb-strain pair and is color-coded by epitope specificity. (**C**) Correlation between Env binding and neutralization. (**D**) Correlation between ADCP and ADCC. Pearson correlations were performed and r and P values are indicated. Nonlinear fit of each correlation is represented by a black line.

We further examined the relationships among bNAb functions within each epitope class (**Figure S12**). V1/V2 apex- and V3 glycan-targeting bNAbs displayed strong positive correlations between Env binding and effector functions, as well as between ADCC and ADCP (**Figure S12B, C**). In contrast, CD4bs-targeting bNAbs mediated effector functions with comparable efficiency regardless of Env binding levels (**Figure S12D**). Similarly, the ability of gp120/gp41 interface-targeting bNAbs and of MPER-targeting bNAbs to trigger effector functions appeared less dependent on binding strength (**Figure S12E, F**). These results suggest that the dependence of effector function on Env binding varied according to the targeted epitope. Notably, neutralization potency positively correlated with both ADCC (r = −0.360, P < 0.0001) and ADCP (r = −0.392, P < 0.0001), demonstrating that bNAbs with stronger neutralization activity were also more effective at mediating Fc-dependent functions (**Figure S12A**).

### Fc engineering enhanced N49P7-mediated effector functions across diverse HIV-1 Envs

Despite moderate Env binding among CD4bs-targeting bNAbs, N49P7 exhibited relatively weak Fc-mediated effector functions, particularly in ADCP. To enhance its activity, we introduced a panel of Fc mutations previously shown to improve ADCC or ADCP in cancer models: ALE (G236A, A330L, I332E), ADE (G236A, S239D, I332E), LPLIL (F243L, R292P, Y300L, V305I, P396L), and DLE (S239D, A330L, I332E) (*44*). To assess the dependence of effector functions on FcγR engagement, we also tested the YTE mutation (M252Y, S254T, T256E), which impairs FcγR binding (*45*). Additionally, we generated an IgG3 version of N49P7 (N49P7-IgG3) by replacing its Fc domain with that of IgG3, a subclass encoding an extended hinge region associated with enhanced effector functions (*18, 40*). To ensure these modifications did not impact antigen interaction, we assessed neutralization potency and Env binding of the variants. Both assays showed comparable results among all N49P7 variants, indicating that Fc modifications did not compromise antigen binding or neutralization activity (**Figure S13, S14**).

Across all strains, the ALE, ADE, LPLIL, and DLE variants consistently enhanced ADCC activity, while YTE and IgG3 decreased this function against some of the strains (**Figure 5A, S15**). In the ADCP assay, the IgG3 variant consistently improved phagocytic activity across most tested strains, whereas the YTE variant impaired it. Mutations such as ADE, LPLIL, and DLE conferred modest ADCP enhancements (**Figure 5B, S16**). Notably, wild-type N49P7 showed little to no ADCC activity against cells expressing HIV_398F1_ Env, HIV_246F3_ Env, and HIV_X1632_ Env, but this could be restored upon introduction of Fc-enhancing mutations (**Figure 5C**). Interestingly, at high concentrations, the YTE and IgG3 variants reduced ADCC activity against HIV_REJO.c_ and HIV_JR-CSF_ Envs (**Figure S15**). Importantly, ADCP activity against Envs that were poorly responsive to wild-type N49P7, such as Envs of HIV_CH119_, HIV_25710_, and HIV_CE1176_, was rescued by IgG3 substitution (**Figure 5D**).

**Figure 5.**
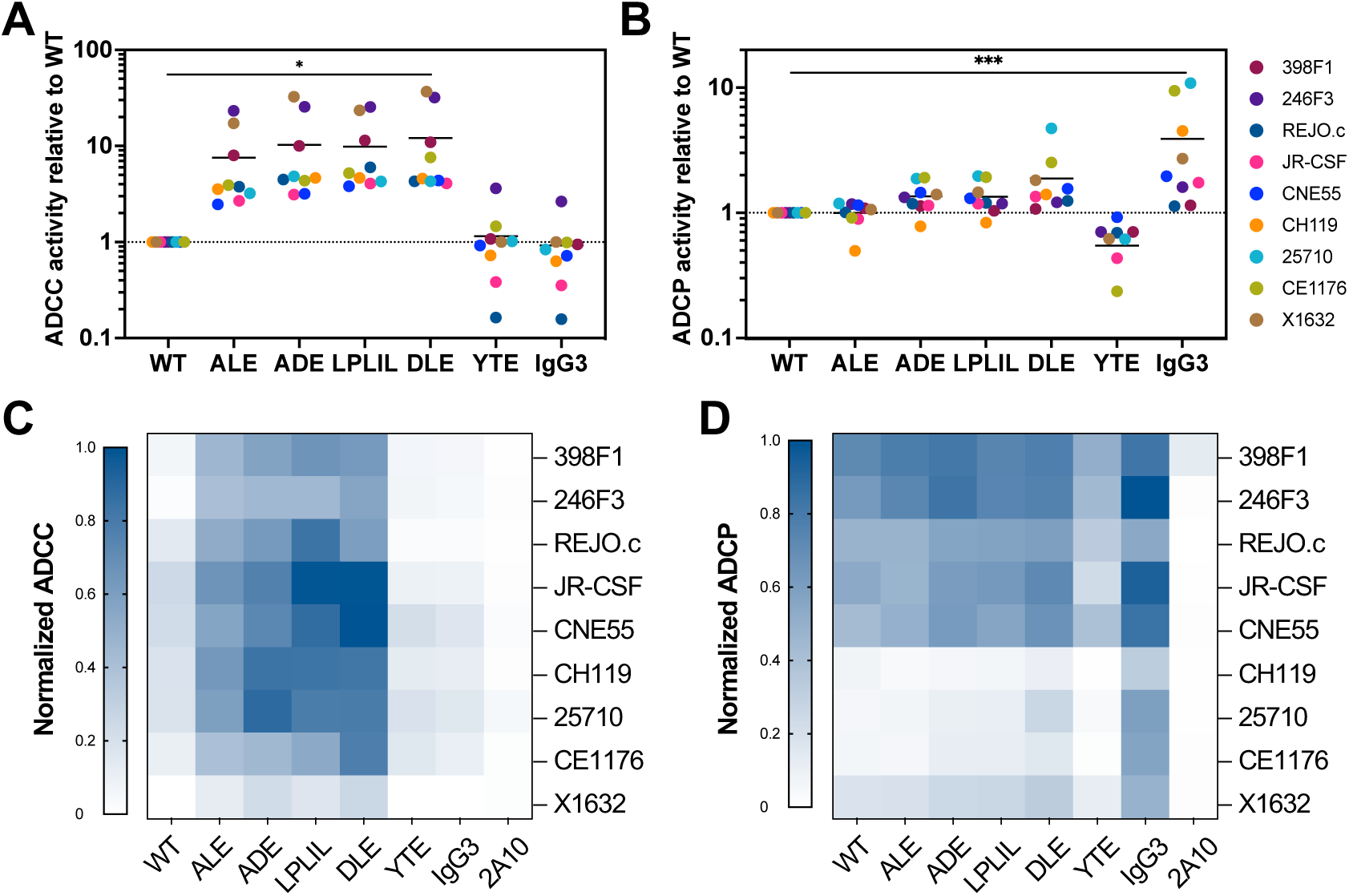
Fc engineering enhances effector functions of N49P7 across diverse strains. Functional characterization of wild-type, Fc-engineered, or IgG3-switched forms of N49P7 against diverse HIV-1 Envs. (**A** and **B**) Normalized ADCC (A) and ADCP (B) activities of Fc-engineered variants relative to wild-type N49P7. Values greater than 1 indicate enhanced activity; values less than 1 indicate reduced activity. (**C** and **D**) Heatmaps showing normalized ADCC (C) and ADCP (D) activities, scaled from 0 to 1 within each effector function across diverse Envs. Fc variants tested include ALE (G236A, A330L, I332E), ADE (G236A, S239D, I332E), LPLIL (F243L, R292P, Y300L, V305I, P396L), DLE (S239D, A330L, I332E), YTE (M252Y, S254T, T256E), and IgG3 (constant region replaced with IgG3).

To assess whether these Fc engineering strategies are generalizable beyond N49P7, we applied the LPLIL, DLE, and YTE mutations, as well as IgG3 Fc substitution, to PGT121, a bNAb targeting the V3 glycan epitope. Fc-engineered PGT121 variants were evaluated for functional activity against HIV_JR-CSF_ Env. Neutralization potency remained comparable against replication-competent HIV_JR-CSF_ across all variants, indicating that Fc modifications did not compromise antigen recognition by PGT121 (**Figure S17A**). Consistent with observations from N49P7, the LPLIL and DLE variants significantly enhanced ADCC relative to wild-type PGT121, whereas the YTE and IgG3 variants reduced ADCC activity (**Figure S17B**). In ADCP assays, the IgG3 variant conferred the greatest enhancement, while DLE and LPLIL provided modest improvements. As seen with N49P7, the YTE mutation reduced ADCP capacity (**Figure S17C**). Notably, all PGT121 variants lost neutralizing activity at antibody concentrations below 0.01 µg/mL and the wild-type PGT121 exhibited minimal ADCC, whereas PGT121 variants carrying the LPLIL or DLE mutation retained ADCC activity. The ADCP activity of wild-type PGT121, PGT121-LPLIL, PGT121-DLE, and PGT121-IgG3 remained potent at low antibody concentrations (**Figure S17**).

Together, these findings show that Fc modifications—particularly ALE, ADE, LPLIL, and DLE—consistently enhanced ADCC across the HIV-1 Envs. However, their effects on ADCP were modest and varied depending on the target Env. In contrast, Fc domain replacement with IgG3 improved ADCP but compromised ADCC. These results demonstrate that rational Fc engineering can selectively tune antibody effector functions and may support strategies to optimize therapeutic efficacy..

## DISCUSSION

Previous studies have shown that antibodies targeting different epitopes exhibit varying levels of ADCC against cells infected with various clade B HIV isolates (*46*). However, most prior studies have focused predominantly on ADCC, with ADCP remaining less explored, which we found to be critical for HIV prevention in humanized mice (*21*). To explore both effector functions across diverse strains from different clades, we generated nine target cell lines stably expressing high levels of HIV Envs at the cell surface. Through systematic profiling of 20 broadly neutralizing antibodies (bNAbs) targeting five distinct epitopes on HIV-1 Env across these diverse target cells, we found that both ADCC and ADCP were shaped by epitope specificity as well as Env sequence diversity. In agreement with prior studies, our analysis showed that CD4bs- and V3 glycan-targeting bNAbs exhibited the most potent effector functions (*29*). Notably, CD4bs-directed bNAbs consistently mediated both ADCC and ADCP across all strains, while V3 glycan-targeting bNAbs displayed potent effector activity against a subset of strains. In contrast, V1/V2 apex-targeting bNAbs primarily induced ADCP, and MPER-targeting bNAbs preferentially mediated ADCC. BNAbs targeting the gp120/gp41 interface showed minimal effector activity across all strains tested (**Figure 1, 2**). These results agree with studies in cancer immunotherapy, which suggest that the location of the epitope relative to the membrane influences effector responses, with membrane-proximal epitopes favoring ADCC and membrane-distal epitopes supporting ADCP (*47*).

In addition to epitope class, our data shows that bNAbs do not always have broad innate functionality, given that diverse Envs exhibited a wide range of functional activity. Individual strains displayed varying sensitivity to ADCC and ADCP. For example, cells expressing HIV_X1632_ Env (clade G) were largely resistant to ADCC, while HIV_CH119_ Env (clade CRF07_BC) showed low ADCP susceptibility across all bNAbs. In contrast, HIV_JR-CSF_ (clade B) and HIV_25710_ (clade C) Envs were highly sensitive to both effector functions across most bNAbs (**Figure 3**). Notably, this variability was not clade-dependent, suggesting that the Env sequence and structure, rather than clade, was important. These results are in agreement with previous studies which found that HIV-infected primary cells exhibited strain-specific differences in ADCC (*29*). Together these findings support earlier reports that structural differences in Env can influence epitope exposure and immune synapse formation, thereby modulating effector functions (*29*).

Interestingly, we observed only modest correlation between Env binding and Fc-mediated functions. For instance, gp120/gp41 interface-targeting bNAbs bound Env at moderate levels but elicited little ADCC or ADCP (**Figure 4**). Even among antibodies targeting the same epitope, function could vary; for example, PG9 and PG16 both target the V1/V2 apex and share overlapping epitopes (*48*), yet PG9 more consistently mediates ADCP across strain as compared to PG16 (**Figure 3**). This is despite equivalent binding activity across these strains, indicating that Env binding alone is not sufficient to predict effector activity. Previous studies suggest that factors such as antibody orientation and stoichiometry of binding likely contribute (*28, 49*). For example, differences in antigen cross-linking and Fc orientation that reduce FcγR engagement have been shown to result in differences in ADCC potency, as demonstrated by experiments in which variable chains were swapped (*28*). These structural and spatial determinants are essential for forming productive immune synapses that are required to trigger downstream effector cascades (*49, 50*).

Conversely, we identified some instances where bNAbs exhibited weak Env binding and neutralization but still mediated effective ADCC and/or ADCP. For example, PGT128 failed to neutralize HIV_REJO.c_ and HIV_CNE55_ and showed relatively lower Env binding, but can trigger effector functions (**Figure 3**). This observation is consistent with previous findings that FcγR triggering can tolerate low-affinity antigen interactions compared to neutralization (*51*). Once activated, NK cells can serially kill multiple infected targets using cytotoxic granules, allowing even modest antibody binding to trigger substantial effector responses (*52*).

Antibody engineering has been extensively employed to enhance Fc-mediated effector function (*53*). For instance, the GASDALIE mutation introduced into 3BNC117 significantly increased its *in vivo* potency in a humanized mouse model (*54*). Therefore, there has been strong interest in Fc-enhancing bNAbs for HIV, including the first-in-class Fc-enhanced bNAb Elipovimab, which is derived from PGT121 and was recently tested in humans (*55, 56*). To enhance bNAb-mediated effector activity, we applied Fc engineering to two representative bNAbs—N49P7 (targeting the CD4 binding site) and PGT121 (targeting the V3 glycan)—using mutations previously shown to increase FcγR binding affinity (44). Variants with ALE, ADE, LPLIL, or DLE substitutions significantly enhanced ADCC across all Envs and bNAbs tested, while ADCP improvements were more modest. The robust enhancement of ADCC is likely due to the increased FcγRIIIa affinity which may improve the serial killing capacity of NK cells of engineered variants (*44*). In contrast, IgG3 substitution increased ADCP but not ADCC, highlighting the distinct roles of FcγRIa/IIa and FcγRIIIa in engaging phagocytes versus NK cells, respectively (**Figure 5**). Notably, introducing specific Fc-region mutations can preserve Fc-mediated effector functions, such as ADCC, even in the absence of neutralizing activity (**Figure S17**). These results suggest that Fc architecture can be rationally tuned to favor specific effector pathways.

Engineered N49P7 and PGT121 variants in our study showed robust ADCC enhancement, improving their activity for therapeutic strategies aimed at eliminating infected cells, such as “shock-and-kill” approaches (*57, 58*). Current latency reversal agents (LRA) aim to reactivate the HIV reservoir, but effective clearance remains a challenge (*59, 60*). Fc-optimized antibodies can enhance the “kill” step by improving functions of innate effector cells. Enhancing ADCP, although more challenging, also remains valuable given previous correlations between ADCP and reduced HIV viremia in NHP model (*61*). From a vaccine design perspective, adjuvants (*62*) and immunogens (*63, 64*) should be selected not only to elicit neutralizing antibodies but also to support favorable Fc profiles. For instance, vaccines targeting CD4bs or V3 glycan epitopes and promoting IgG subclasses (*21*) or glycoforms (*27*) that efficiently engage FcγRIIa and FcγRIIIa may enhance vaccine-induced protection.

Our study has a number of limitations. For instance, all functional assays were conducted *in vitro* using immortalized cell lines, which may not fully recapitulate *in vivo* immune responses. For instance, NK-92 cells lack the receptor diversity and education observed in primary NK cells (*65, 66*), and THP-1 cells may not fully replicate the complex behaviors and accurately reflect the *in vivo* conditions exhibited by primary blood monocytes (*67*). Additionally, Env expression on engineered cells exceeds physiological levels due to a lack of regulation seen in infected cells, which often show low Env density (*68*) or control Env conformation (*69*) to limit effector function engagement. Additionally, while we observed only minor differences in Env expression across strains (**Figure S7B**), it is possible that these differences also contributed to differential activity measured in our assays. Moreover, HIV infection can evade immune clearance by downregulating NK-activating ligands (*70*) or antagonizing tetherin to prevent antibody-dependent cytotoxicity (*71*), which is not present in our model. These caveats highlight the need for *in vivo* validation, including studies in humanized mouse or nonhuman primate models, to assess efficacy, pharmacokinetics, immunogenicity, and escape potential.

In summary, our findings show that bNAb-mediated effector functions are governed by a complex interplay between epitope specificity, Env sequence and structure, and Fc architecture. Rational Fc engineering could substantially augment ADCC and ADCP across genetically diverse HIV strains and offer improved activity against strains that are poorly neutralized. These insights highlight the importance of testing and optimizing bNAbs across strains to ensure that broad neutralizing activity is associated with equally broad innate functionality.

## MATERIALS AND METHODS

### Study design

This study aimed to investigate the antibody-mediated effector functions of HIV-1 bNAbs across diverse Env variants. A panel of twenty bNAbs was evaluated for neutralization, Env binding, ADCC, and ADCP using replication-competent virus or Env-expressing cell models. Each experiment was independently repeated at least three times to ensure reproducibility.

### Cell lines

HEK293T cells were obtained from the American TypeCulture Collection and TZM-bl Cells (ARP-8129) were obtained through the NIH HIV Reagent Program, Division of AIDS, NIAID, NIH, contributed by John C. Kappes, Xiaoyun Wu and Tranzyme Inc. The HEK293T and TZM-bl cells were cultured in DMEM (Corning) containing 10% fetal bovine serum (Millipore Sigma) and Penicillin-Streptomycin (Fisher Scientific) at 37°C/5% CO_2_. CEM.NKr cells were obtained from the NIH AIDS Reagent Program and engineered to express HIV Env using lentiviral transduction. CEM.NKr and Env-expressing CEM.NKr cells were cultured in RPMI 1640 (Stemcell) containing 10% fetal bovine serum (Millipore Sigma), 2mM L-glutamine (Life Technologies), and Penicillin-Streptomycin (Fisher Scientific) at 37°C/5% CO_2_. THP-1 cells were obtained from the American TypeCulture Collection and were cultured under the same conditions as CEM.NKr cells. NK-92 cells were obtained from the American TypeCulture Collection and engineered to express CD16 (FcγRIIIa V158), and cultured in RPMI 1640 containing 10% fetal bovine serum, 2mM L-glutamine, Penicillin-Streptomycin, and 0.01ng/mL IL-2.

### Construction and purification of broadly neutralizing antibody panel

The coding sequences for the variable and constant regions of both heavy and light chains of each bNAb were cloned into an expression vector. A self-cleaving 2A peptide was inserted between the heavy and light chain coding regions to promote ribosomal skipping during translation, enabling co-expression from a single transcript. All constructs were verified by Sanger sequencing to confirm sequence accuracy. Expression and purification of recombinant bNAbs was through established methodology (*72–74*). Briefly, Expi293 cells were transfected with the expression vector using the ExpiFectamine™ 293 Transfection Kit (Thermofisher), which supplies the transfection reagent and enhancer solution. Six days after transfection, the culture supernatants were harvested, filtered using VacuCap 8/0.2 μm filters (Pall Corporation), and buffer then exchanged into PBS using a tangential flow filtration (TFF) system [Pall Corporation; T-Series Centramate cassettes with Omega PES membrane 10 kDa (Cytiva, OS010T12)]. The bNAbs were purified by affinity chromatography using Pierce Protein A/G Agarose resin (Thermo Fisher Scientific) followed by size-exclusion chromatography using a Superdex 200 column (GE Healthcare) on an ÄKTA Purifier Fast Protein Liquid Chromatography (FPLC) system (GE Healthcare). Antibody concentrations were quantified using a NanoDrop 2000 Spectrophotometer (Thermo Fisher Scientific).

### HIV production and quantification

To generate chimeric HIV-1 strains, the full-length Env gene from HIV_398F1_, HIV_246F3_, HIV_CNE55_, HIV_CH119_, HIV_25710_, HIV_CE1176_, or HIV_X1632_ was cloned into HIV_JR-CSF_ backbone pYK-JRCSF (ARP-2708, obtained from the NIH AIDS Reagent Program) to replace the original Env sequence. This resulted in the following chimeric viruses: HIV_JR-CSF-398F1_, HIV_JR-CSF-246F3_, HIV_246F3-CNE55_, HIV_JR-CSF-CH119_, HIV_JR-CSF-CE1176_, and HIV_JR-CSF-X1632_. Viruses were produced by transient transfection of HEK293T cells with 1 μg/mL of plasmid DNA encoding HIV_REJO.c_ (ARP-11746, pREJO.c/2864, NIH AIDS Reagent Program), HIV_JR-CSF_, or the respective chimeric constructs. After 48 hours, supernatants were collected, filtered through a 0.45 μm filter, and quantified using a 50% tissue culture infective dose (TCID50) assay on TZM-bl cells. TCID_50_ values were calculated using the Spearman-Karber formula (*75*).

### Neutralization assay

Neutralization assays were performed using a Fluent Automated Workstation (Tecan) liquid handler in 384-well plates (Greiner) and conducted in quadruplicate. Each well received 200 TCID_50_ of virus combined with a 2.5-fold serial dilution of the test antibody. The virus-antibody mixtures were incubated at room temperature for 1 hour, followed by the addition of 6,000 TZM-bl cells per well. Cultures were incubated at 37°C with 5% CO₂ for 48 hours in the presence of 75 μg/mL DEAE-dextran to enhance viral entry. After incubation, cells were lysed using a previously described luciferase lysis buffer (*76*), and luminescence was measured using a SpectraMax L microplate reader (Molecular Devices). Percentage of infection was determined by calculating the difference in luminescence between experimental wells (cells with virus and antibody) and cell control wells (cells only) and dividing this value by the difference between the virus control wells (cells with virus) and the cell control wells. Percent neutralization was determined by subtracting the infection from 100%. Neutralization curves were generated by plotting percent neutralization against antibody concentration, and IC_50_ values were calculated using a three-parameter nonlinear regression in GraphPad Prism 10.0.

### Generation of HIV Env-Expressing CEM.NKr Cells

CEM.NKr cells were obtained from the NIH AIDS Reagent Program and engineered to express membrane-bound HIV-1 gp150 from HIV_398F1_, HIV_246F3_, HIV_REJO.c_, HIV_JR-CSF_, HIV_CNE55_, HIV_CH119_, HIV_25710_, HIV_CE1176_, or HIV_X1632_ using lentiviral transduction as previously mentioned (21). Briefly, the gene of HIV gp150 was codon-optimized for enhanced expression and modified by replacing the endogenous leader peptide with the signal sequence of human CD5 antigen (*77*) and truncating the C-terminus to remove endocytosis signals (*21*). The resulting gene was synthesized (Integrated DNA Technologies) and cloned into a lentiviral expression vector (*78*) under the control of the human EF1α promoter. An internal ribosome entry site (IRES) positioned downstream of the Env gene facilitated cap-independent translation of ZsGreen, a fluorescent protein used for identifying successfully transduced cells. Lentiviral particles were generated by co-transfecting HEK293T cells with the gp150 lentiviral plasmid and four helper plasmids (pHDM-VSV-G, pHDM-Hgpm2, pHDM-Tat1b, and pRC-CMV-Rev1b). After 48 hours, viral supernatants were harvested, filtered through a 0.45 µm filter, and used to transduce CEM.NKr cells. Following transduction, CEM.NKr cells were bulk sorted using a FACSAria 4II SORP Flow Cytometer (BD Biosciences). Single-cell clones were then generated by limiting dilution and screened for high expression of both HIV Env and ZsGreen fluorescence to establish stably transduced target cell lines for downstream functional assays.

### Cell-associated envelope binding assay

To assess antibody binding to cell-surface HIV-1 Env, antibodies were conjugated with Alexa Fluor 647 (AF647) using the Zip Alexa Fluor™ 647 Rapid Antibody Labeling Kit (Thermo Fisher Scientific) according to the manufacturer’s instructions. Env-expressing CEM.NKr cells were mixed at a 1:1 ratio with parental CEM.NKr cells, which serve as an internal control for background staining. The mixed cell population was incubated with three-fold serial dilutions of each AF647-labeled antibody for 30 minutes at room temperature. After incubation, cells were washed twice with PBS containing 2% fetal bovine serum (PBS+), fixed with 4% paraformaldehyde (Fisher Scientific), and analyzed on a Stratedigm S1300Exi flow cytometer. Median fluorescence intensity (MFI) of the AF647 signal was calculated for each antibody to quantify Env-specific binding.

### Antibody-dependent cellular cytotoxicity (ADCC) assay

Env-expressing CEM.NKr cells were mixed with parental CEM.NKr cells at a 1:1 ratio. The mixed target population (5 x 10^3^ Env-expressing CEM.NKr and 5 x 10^3^ parental CEM.NKr cells per well) was incubated with three- or four-fold serial dilutions of antibody, starting at 30 µg/mL or 2.5 µg/mL, for 20 minutes at room temperature. CD16.NK-92 cells (NK-92 cells stably expressing CD16) were stained with CellTrace Violet (Thermo Fisher Scientific) for 20 minutes at room temperature and added to the antibody-target cell mixture at a 5:1 effector-to-target ratio (5 x 10^4^ effector cells per well). Plates were incubated at 37 °C for 6 hours. Following incubation, cells were washed with PBS+, fixed with 4% paraformaldehyde, and then analyzed on a Stratedigm S1300Exi flow cytometer. Target cell lysis was calculated by subtracting the percentage of ZsGreen⁺ Env-expressing CEM.NKr cells in the experimental wells (with effector cells and antibody) from the percentage in the control wells (with effector cells only), and dividing this difference by the percentage in the control wells.

### Antibody-dependent cellular phagocytosis (ADCP) assay

ZsGreen⁺ Env-expressing CEM.NKr target cells were incubated with three- or four-fold serial dilutions of antibody, starting at 10 µg/mL or 0.25 µg/mL, for 20 minutes at room temperature. After incubation, THP-1 effector cells labeled with CellTrace Violet were added at a 2:1 effector-to-target ratio (5 x 10^4^ THP-1 cells and 2.5 x 10^4^ target cells per well). Plates were incubated at 37 °C for 1 hour. Following incubation, cells were fixed with 4% paraformaldehyde for 20 minutes and analyzed on a Stratedigm S1300Exi flow cytometer. Phagocytosis was quantified as the percentage of THP-1 cells with ZsGreen signal.

### Construction and purification of Fc variant antibodies

Fc variants were generated by introducing point mutations into the wild-type antibody transgene via PCR with mutation-specific primers or by substituting the IgG1 constant region with the IgG3 sequence. Modified constructs were cloned into an expression vector. All constructs were verified by Sanger sequencing to confirm sequence accuracy and were expressed and affinity purified as described above.

### Quantification and statistical analysis

Data and statistical analyses were performed using GraphPad Prism v10.0.3. Flow cytometry data was analyzed using FlowJo v10.8.2. For all figures statistical significance was defined by GraphPad Prism as * = p < 0.0332, ** = p < 0.0021, *** = p < 0.0002, **** = p < 0.0001. Phylogenetic Tree was built using Geneious Prime v2023.0.4.

## Acknowledgments

We wish to thank members of the Balazs lab for their helpful suggestions and discussions.

## Funding

Postdoctoral Research Abroad Program from the National Science and Technology

Council (NSTC), Taiwan (Grand No. NSTC-113-2917-I-564-017) to C.J.L.

National Institute of Allergy and Infectious Diseases (NIAID) R01AI174875 to A.B.B.

National Institute of Allergy and Infectious Diseases (NIAID) R01AI174276 to A.B.B.

National Institute on Drug Abuse (NIDA) DP1DA060607 to A.B.B.

National Institute on Drug Abuse (NIDA) DP2DA040254 to A.B.B.

National Institute of Allergy and Infectious Diseases (NIAID) R01AI155447 to D.L.

National Institute of Allergy and Infectious Diseases (NIAID) R01AI137057 to D.L.

National Institute of Allergy and Infectious Diseases (NIAID) R01AI153098 to D.L.

National Institute of Allergy and Infectious Diseases (NIAID) P30AI060354 to D.L.

## Author contributions

Conceptualization: C.J.L., M.P., J.M.B., and A.B.B.

Methodology: C.J.L., M.P., J.M.B., V.O., C.M., C.A., M.E.A., D.L., and A.B.B.

Investigation: C.J.L., E.L., and M.P.

Visualization: C.J.L., E.L., and M.P.

Funding acquisition: C.J.L,. D.L., and A.B.B.

Writing – original draft: C.J.L., M.P., and A.B.B.

Writing – review & editing: C.J.L., E.L, M.E.A. and A.B.B.

Supervision: C.J.L., D.L., and A.B.B.

## Data and materials availability

Cell lines and replication-competent chimeric viruses are deposited with BEI resources. Lentiviral vectors expressing diverse HIV-1 Envs are deposited with Addgene. This study did not generate sequence data or code. Raw data generated in the current study (including Env binding, ADCC, ADCP, and neutralization assay) have not been deposited in a public repository but are available from the corresponding author upon request. Plasmids generated in this study are available from A.B.B. under material transfer agreement with MGH.

