## Supplemental Figures for "Functional Activity of HIV-1 bNAbs Across Diverse Strains is Driven by Binding Site and Can be Enhanced Through Fc Engineering"

Table S1. Panel of broadly neutralizing antibodies (bNAbs).

| **Epitope** | V1/V2 apex | V3 glycan | CD4bs | gp120/gp41 interface | MPER |
|---|---|---|---|---|---|
| **Antibody** | PG9<br>PG16<br>PGDM1400 | PGT121<br>PGT128<br>10-1074 | VRC07<br>N6<br>N49P7<br>VRC13<br>VRC01.23LS<br>VRC07-523LS<br>1-18<br>3BNC117 | 35O22<br>PGT151<br>8ANC195 | PGZL1.H4K3<br>10E8v4-V5R-100cF<br>4E10 |

Table S2. Panel of HIV-1 strains.

| **Isolate Original Name** | **Isolate Common Name** | **Clade** |
|---|---|---|
| 398_F1_F6_20 | 398F1 | A |
| 246_F3_C10_2 | 246F3 | AC |
| REJO.c | REJO.c | B |
| JR-CSF | JR-CSF | B |
| CNE55 | CNE55 | CRF01_AE |
| CH119.10 | CH119 | CRF07_BC |
| HIV_ 25710-2.43 | 25710 | C |
| CE1176_A3 | CE1176 | C |
| X1632_S2_B10 | X1632 | G |

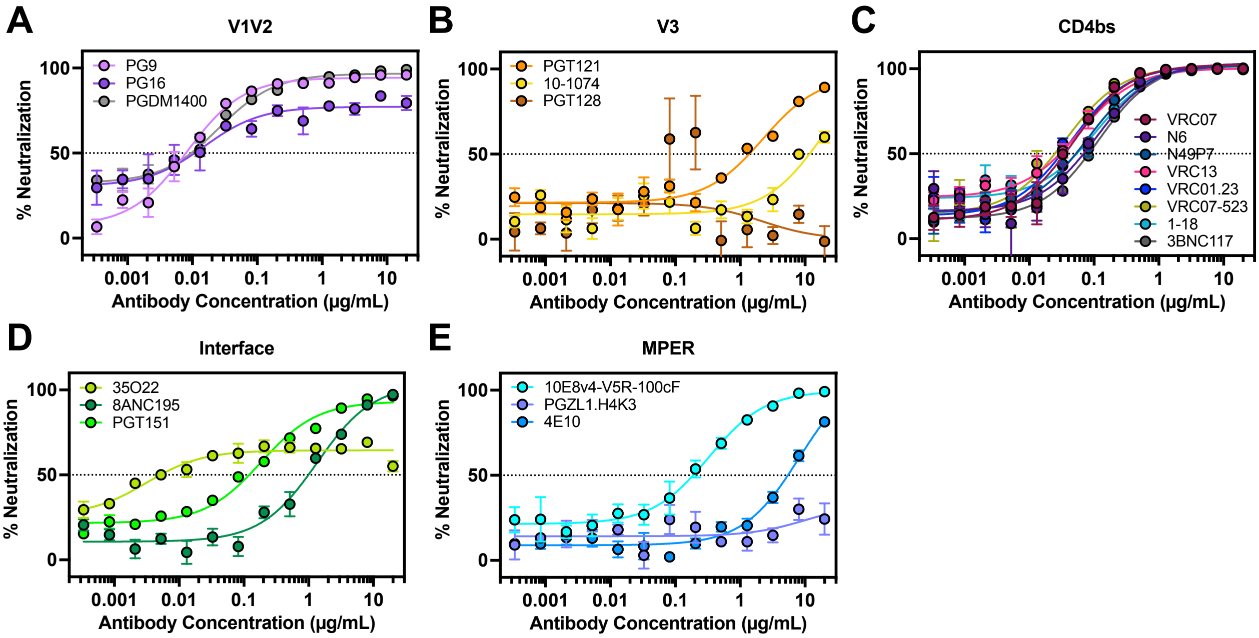


**Figure S1. Neutralization assays of HIV-1 bNAbs against HIV$_{REJO.c}$.**
(**A-E**) Neutralization activity of bNAbs was assessed against replication-competent HIV$_{REJO.c}$. Data are shown as percent neutralization across a range of antibody concentrations. Antibodies are grouped by epitope: V1/V2 apex (A), V3 glycan (B), CD4 binding site (C), gp120/gp41 interface (D), and MPER (E), with individual bNAbs indicated in the legends. Neutralization assays were performed in quadruplicate.

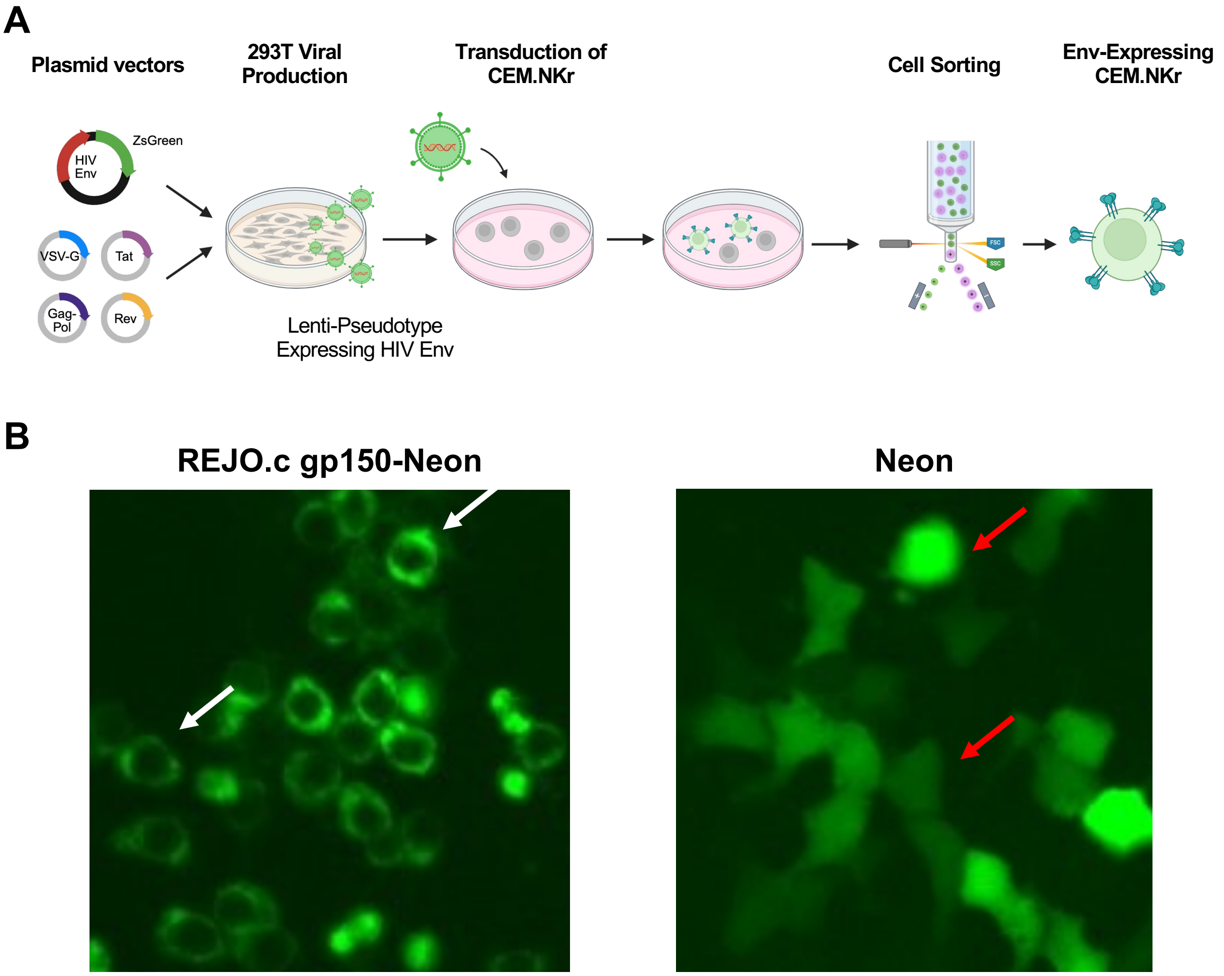


**Figure S2. Generation of HIV Env-expressing CEM.NKr cells.**
(**A**) Schematic overview of the Env-expressing CEM.NKr development. Lentiviral particles were generated by co-transfecting HEK293T cells with Env and ZsGreen-expressing lentiviral plasmid (pHAGE2) and four helper plasmids (pHDM-VSV-G, pHDM-Hgpm2, pHDM-Tat1b, and pRC-CMV-Rev1b). Viral supernatant was collected to transduce the CEM.NKr cells. After 48 hours, CEM.NKr cells were bulk sorted by flow cytometry. Single-cell clones were then generated by limiting dilution and screened for high expression of both HIV Env and ZsGreen fluorescence to establish stably transduced target cell lines. (**B**) HEK293T cells transfected with Neon-tagged $HIV_{REJO.c}$ Env exhibit Env surface expression (white arrows), while cells transfected with Neon alone display cytoplasmic fluorescence (red arrows).

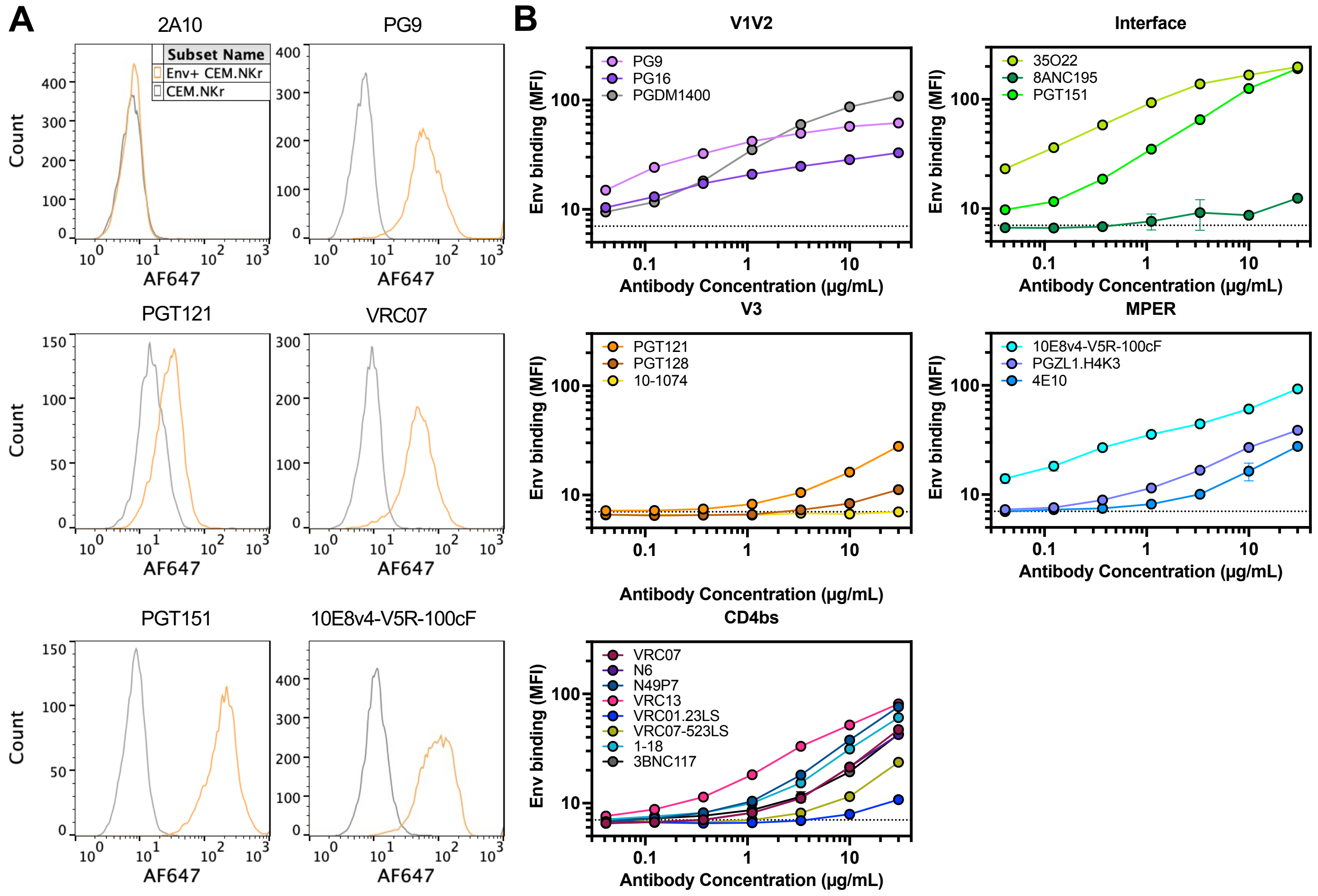


**Figure S3. Env binding of HIV-1 bNAbs to $HIV_{REJO.c}$ Env-expressing CEM.NKr cells.** (**A**) Binding of 30 µg/mL of AF647-conjugated HIV bNAbs to CEM.NKr cells expressing $HIV_{REJO.c}$ Env. The orange curves represent bNAb binding to $HIV_{REJO.c}$ Env-expressing cells, while the gray curves show binding to parental CEM.NKr cells. A malaria-specific IgG1 2A10 was included as control. (**B**) Binding of twenty AF647-conjugated HIV bNAbs to CEM.NKr cells expressing $HIV_{REJO.c}$ Env across a range of bNAb concentrations, with individual antibodies indicated in the legends. Binding is presented as mean fluorescence intensity (MFI) at each concentration and was performed in triplicate. Dashed line indicates control IgG1 2A10 signal.

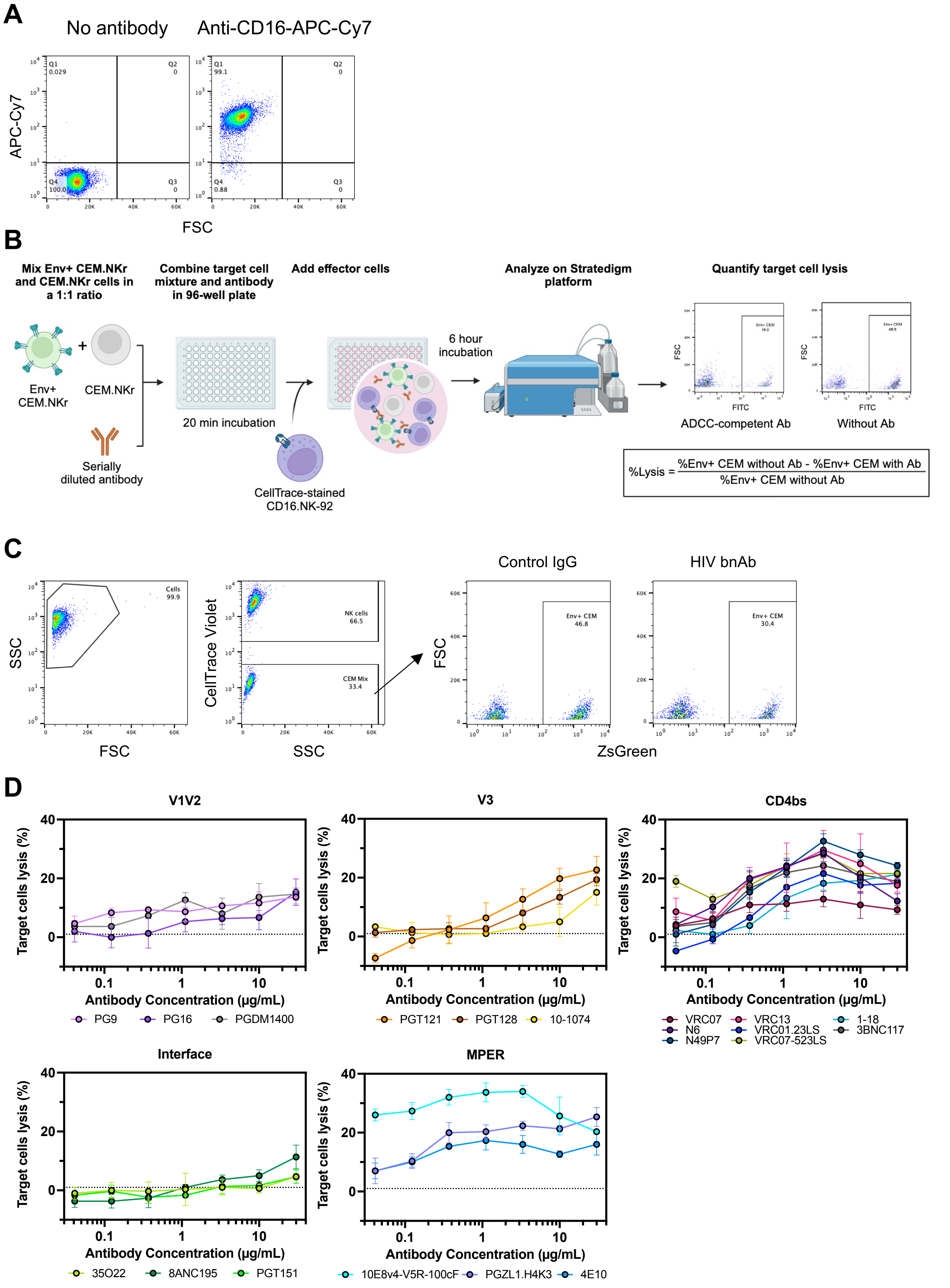
A
No antibody
Anti-CD16-APC-Cy7
APC-Cy7
FSC
Q1 0.029
Q2 0
Q3 0
Q4 100.0
Q1 99.1
Q4 0.88
B
Mix Env+ CEM.NKr and CEM.NKr cells in a 1:1 ratio
Combine target cell mixture and antibody in 96-well plate
Add effector cells
Analyze on Stratedigm platform
Quantify target cell lysis
Env+ CEM.NKr
CEM.NKr
Serially diluted antibody
20 min incubation
CellTrace-stained CD16.NK-92
6 hour incubation
ADCC-competent Ab
Without Ab
%Lysis = (%Env+ CEM without Ab - %Env+ CEM with Ab) / %Env+ CEM without Ab
C
Cells 99.9
SSC
FSC
CellTrace Violet
NK cells 66.5
CEM Mix 33.4
Control IgG
HIV bnAb
Env+ CEM 46.8
Env+ CEM 30.4
ZsGreen
D
V1V2
V3
CD4bs
Interface
MPER
Target cells lysis (%)
Antibody Concentration (µg/mL)
PG9
PG16
PGDM1400
PGT121
PGT128
10-1074
VRC07
N6
N49P7
VRC13
VRC01.23LS
VRC07-523LS
1-18
3BNC117
35O22
8ANC195
PGT151
10E8v4-V5R-100cF
PGZL1.H4K3
4E10

**Figure S4. Development of CD16.NK-92 effector cells and flow cytometry-based ADCC assay.**

(**A**) Generation and validation of CD16-expressing NK-92 cells (CD16.NK-92). Flow cytometry analysis confirmed surface expression of CD16. (**B**) Schematic overview of the flow cytometry-based ADCC assay. ZsGreen+ Env-expressing CEM.NKr target cells were mixed with parental CEM.NKr cells and incubated with serially diluted antibodies. CellTrace Violet-labeled CD16.NK-92 effector cells were then added, followed by a 6-hour incubation. After co-culture, cells were analyzed by flow cytometry to quantify target cell lysis using the formula shown. (**C**) Representative flow cytometry gating strategy. Cells were first gated on FSC and SSC, followed by separation of effector (CellTrace$^{+}$) and target (CellTrace$^{-}$) populations. ZsGreen$^{+}$ target cells (Env-expressing CEM.NKr) were quantified in the presence of control IgG or HIV bNAbs. A decrease in the proportion of ZsGreen$^{+}$ CEM.NKr cells relative to parental CEM.NKr cells in the bNAb-treated condition indicates ADCC-mediated target cell lysis. (**D**) ADCC of twenty HIV bNAbs against CEM.NKr cells expressing $HIV_{REJO.c}$ Env, with individual bNAbs indicated in the legends. ADCC activity was evaluated across a range of antibody concentrations and was shown as percent target cell lysis. Dashed line indicates control IgG1 2A10 activity.

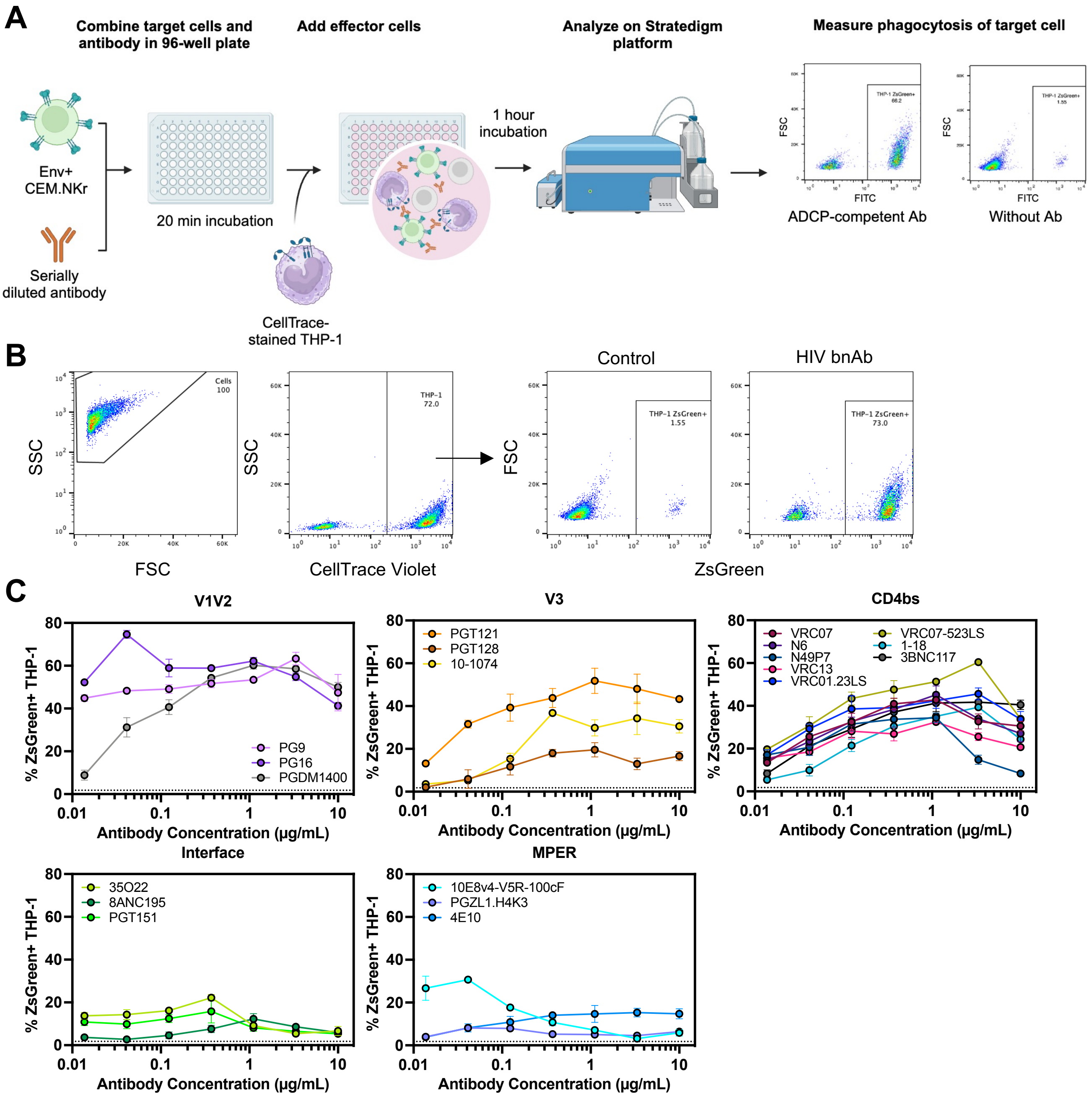


**Figure S5. Flow cytometry-based ADCP assay.**
(**A**) Schematic overview of the flow cytometry-based ADCP assay. ZsGreen+ Env-expressing CEM.NKr target cells were incubated with serially diluted antibodies. CellTrace Violet-labeled THP-1 effector cells were then added, followed by a 1-hour incubation. After co-culture, cells were analyzed by flow cytometry. Phagocytosis was measured as the percentage of THP-1 cells that have taken up ZsGreen$^{+}$ CEM.NKr target cells, indicative of ADCP activity. (**B**) Representative flow cytometry gating strategy. Cells were first gated on FSC and SSC, and THP-1 effector cells were identified based on CellTrace Violet staining. ZsGreen$^{+}$ THP-1 cells were then quantified to assess phagocytic uptake of Env$^{+}$ target cells. Representative plots show minimal uptake in the control IgG condition and substantial uptake in the presence of an HIV bNAb. (**C**) ADCP of twenty HIV bNAbs against CEM.NKr cells expressing $HIV_{REJO.c}$ Env, with individual bNAbs indicated in the legends. ADCP activity was measured across a range of antibody concentrations and was determined by measuring the phagocytosis of ZsGreen+ $HIV_{REJO.c}$ Env-expressing CEM.NKr cells by THP-1 effector cells. Dashed line indicates control IgG1 2A10 activity.

**A**

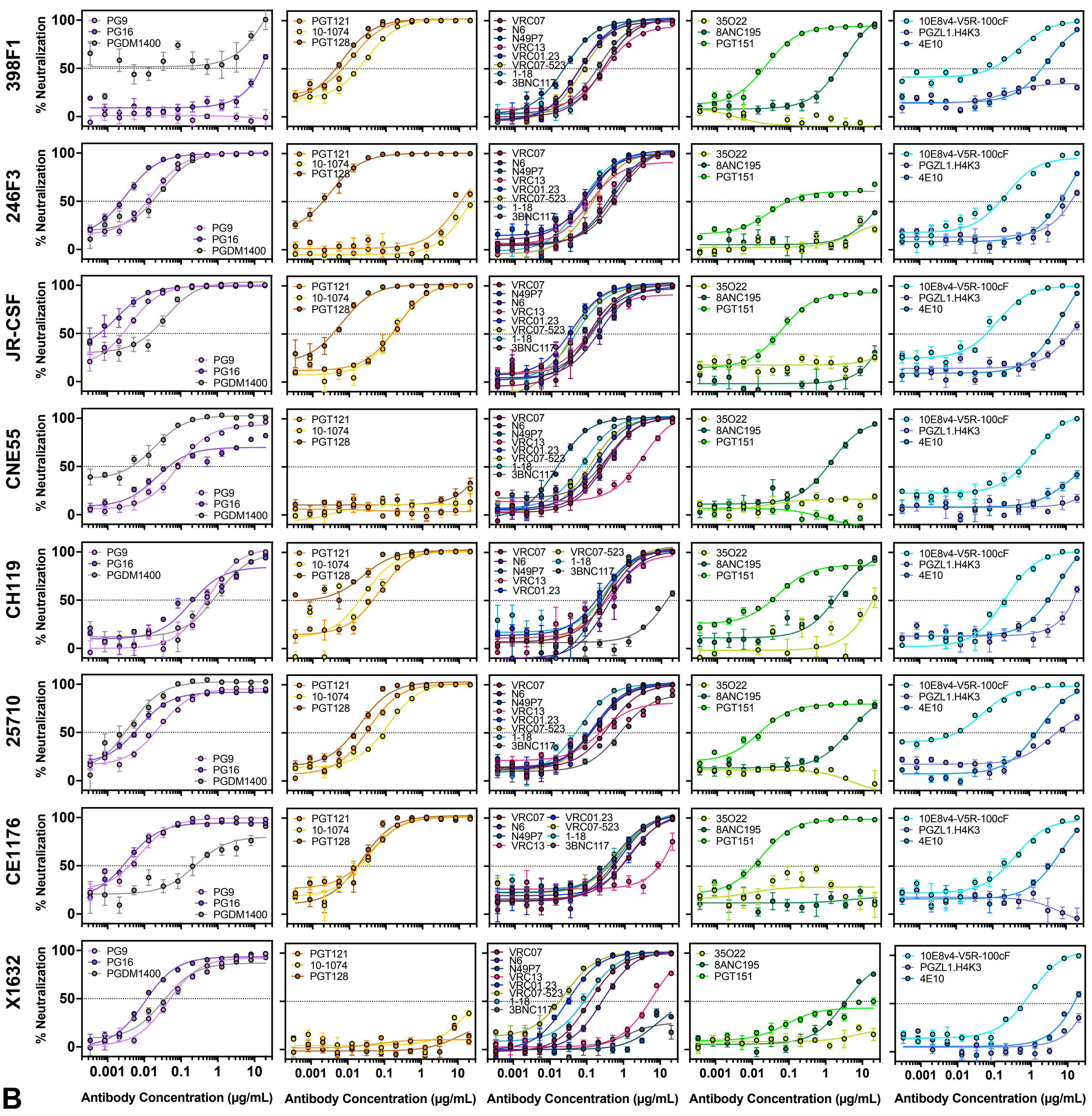


**B**

| | V1V2 | | | V3 | | | CD4bs | | | | | | | | Interface | | | MPER | | | |
|---|---|---|---|---|---|---|---|---|---|---|---|---|---|---|---|---|---|---|---|---|---|
| | PG9 | PG16 | PGDM 1400 | PGT 121 | PGT 128 | 10-1074 | VRC07 | N6 | N49P7 | VRC13 | VRC01 .23LS | VRC07 -523LS | 1-18 | 3BNC 117 | 35O22 | 8ANC 195 | PGT 151 | 10E8v 4-V5R-100cF | PGZL1. H4K3 | 4E10 | 2A10 |
| 398F1 | >20 | >20 | 18.32 | 0.01 | 0.01 | 0.02 | 0.22 | 0.04 | 0.02 | 0.20 | 0.06 | 0.06 | 0.19 | 0.17 | >20 | 2.23 | 0.02 | 0.23 | >20 | 2.74 | >20 |
| 246F3 | 0.01 | 0.00 | 0.02 | 8.70 | 0.00 | >20 | 0.49 | 0.05 | 0.28 | 0.08 | 0.11 | 0.11 | 0.05 | 0.60 | >20 | >20 | 0.02 | 0.04 | 17.11 | 12.37 | >20 |
| REJO.c | 0.01 | 0.00 | 0.02 | 1.33 | >20 | 12.34 | 0.04 | 0.08 | 0.05 | 0.04 | 0.04 | 0.03 | 0.05 | 0.10 | 0.00 | 1.11 | 0.10 | 0.27 | >20 | 8.33 | >20 |
| JR-CSF | 0.00 | 0.00 | 0.03 | 0.18 | 0.00 | 0.20 | 0.11 | 0.14 | 0.08 | 0.07 | 0.04 | 0.03 | 0.04 | 0.14 | >20 | >20 | 0.04 | 0.10 | 12.88 | 7.28 | >20 |
| CNE55 | 0.06 | 0.02 | 0.01 | >20 | >20 | >20 | 0.28 | 0.24 | 0.01 | 1.78 | 0.14 | 0.11 | 0.06 | 0.24 | >20 | 0.95 | >20 | 0.73 | >20 | >20 | >20 |
| CH119 | 0.44 | 0.18 | 0.79 | 0.05 | 0.01 | 0.01 | 0.31 | 0.22 | 0.15 | 0.19 | 0.24 | 0.24 | 0.23 | 15.67 | 0.01 | 1.62 | 0.02 | 0.16 | >20 | 5.62 | >20 |
| 25710 | 0.02 | 0.00 | 0.00 | 0.03 | 0.02 | 0.10 | 0.40 | 0.15 | 0.15 | 0.10 | 0.14 | 0.13 | 0.06 | 0.61 | >20 | 3.55 | 0.01 | 0.03 | 2.85 | 1.45 | >20 |
| CE1176 | 0.01 | 0.00 | 0.12 | 0.04 | 0.02 | 0.03 | 0.79 | 0.64 | 0.45 | >20 | 1.36 | 0.54 | 0.82 | 0.59 | >20 | >20 | 0.01 | 0.41 | >20 | 6.22 | >20 |
| X1632 | 0.04 | 0.01 | 0.02 | >20 | >20 | >20 | 0.11 | 0.26 | >20 | 5.87 | 0.03 | 0.02 | 0.08 | >20 | >20 | 3.39 | >20 | 0.77 | >20 | >20 | >20 |

**Figure S6. Neutralization assay of bNAbs to diverse HIV-1 strains.**
(**A**) Neutralization activity of bNAbs was assessed against diverse HIV-1 strains. Data are shown as percent neutralization across a range of antibody concentrations. Assays were performed in quadruplicate. (**B**) Table summarizing the $IC_{50}$ values for each antibody against each strain.

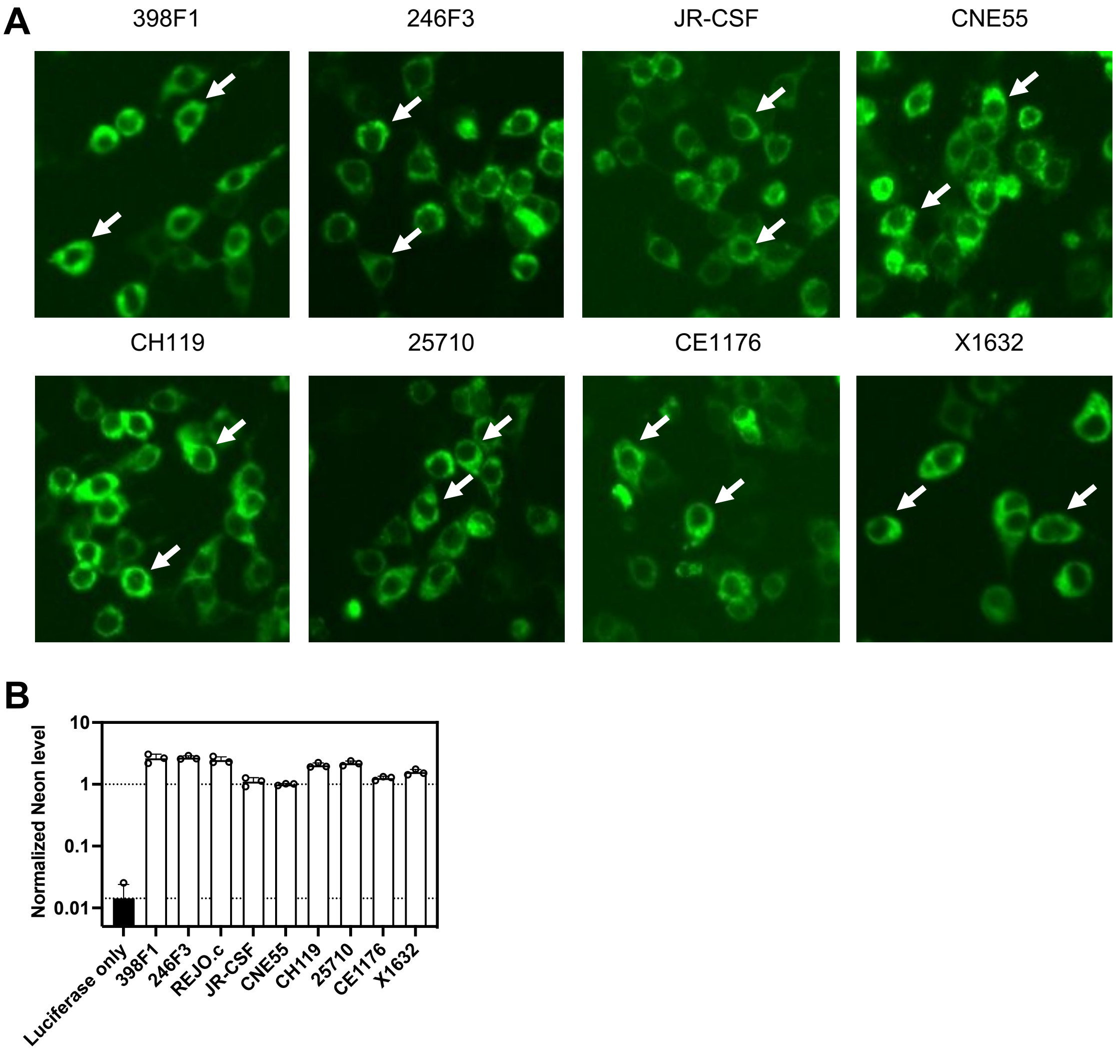


**Figure S7. Surface expression of diverse HIV-1 Envs on transfected cells.**
(A) Fluorescence microscopy images of HEK293T cells transfected with Neon-tagged gp150 constructs. A luciferase plasmid was co-transfected as a control for transfection efficiency. All Env proteins localized to the cell surface, as indicated by peripheral fluorescence signals (white arrows). (B) Mean fluorescence intensity (MFI) of Neon-positive cells was normalized to the corresponding luminescence signal and further normalized to that of CNE55.

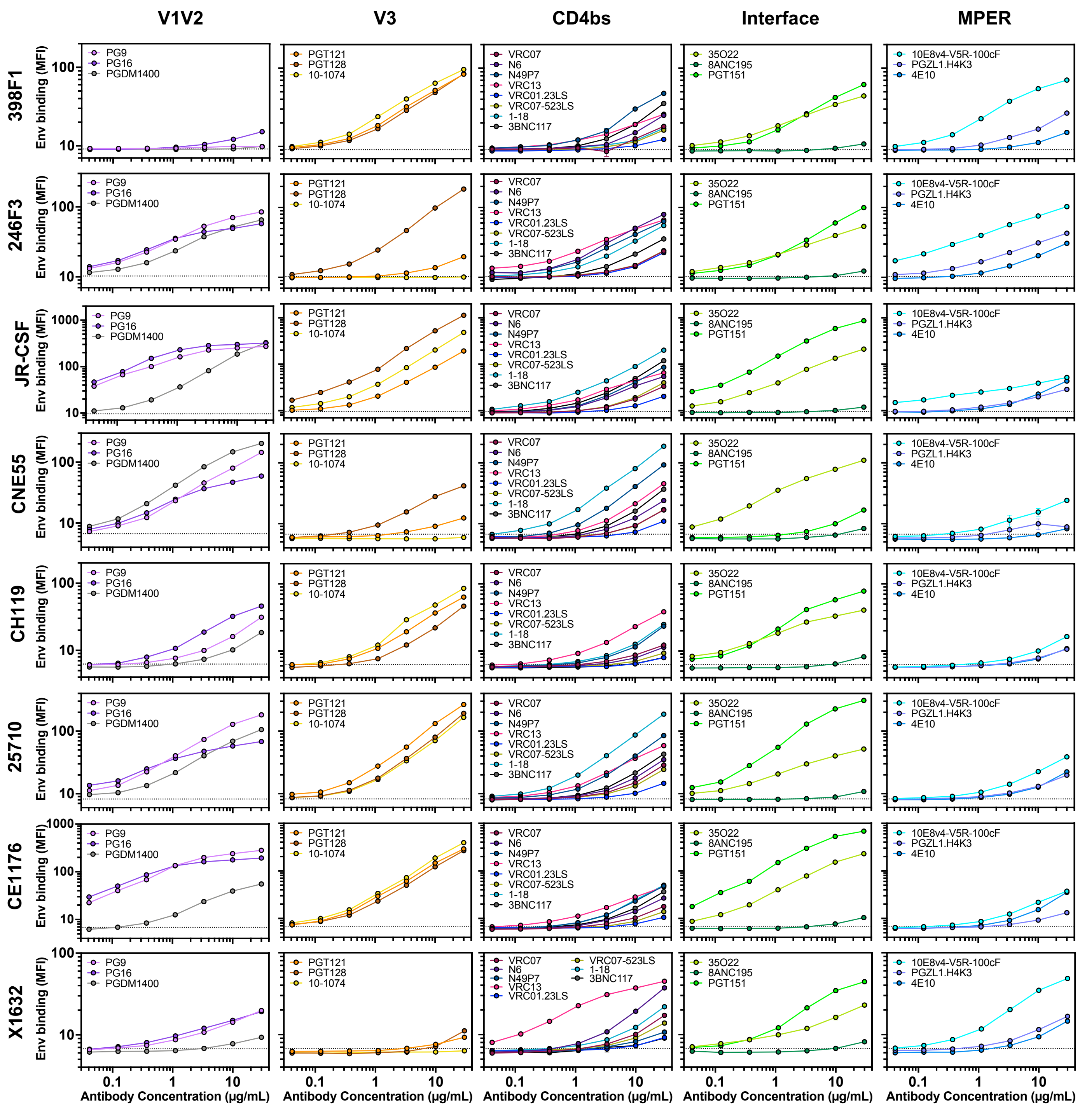


**Figure S8. Env binding of bNAbs to Env-expressing CEM.NKr cells.**
Binding of twenty AF647-conjugated bNAbs to CEM.NKr cells expressing each of the diverse HIV-1 Envs, with individual bNAbs indicated in the legends. Binding is presented as mean fluorescence intensity (MFI) at each bNAb concentration and was performed in triplicate.

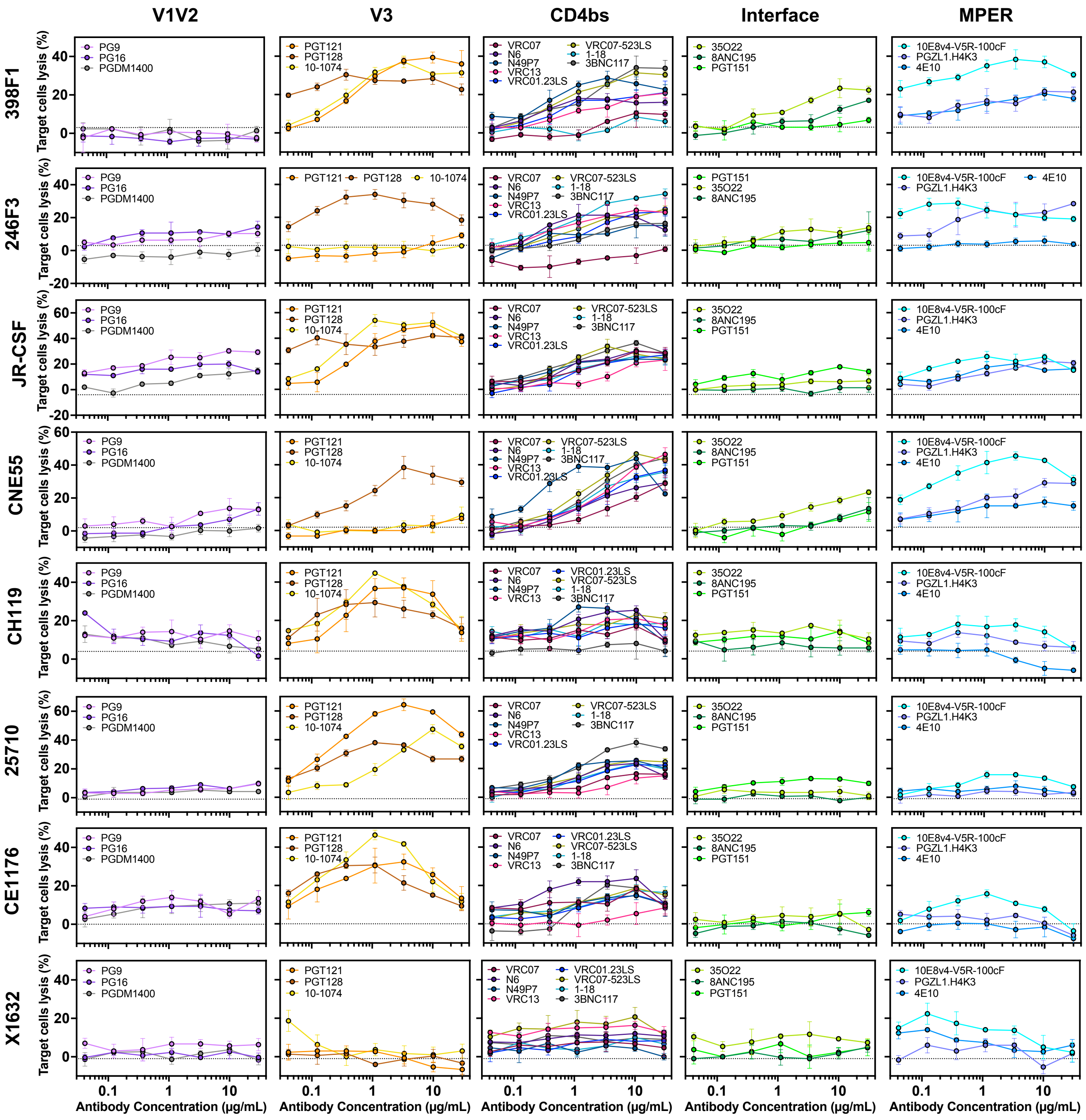


**Figure S9. ADCC of bNAbs to Env-expressing CEM.NKr cells.**
ADCC of bNAbs against CEM.NKr cells expressing each of the diverse HIV-1 Envs, with individual bNAbs indicated in the legends. ADCC activity was performed as mentioned in Fig. S4.

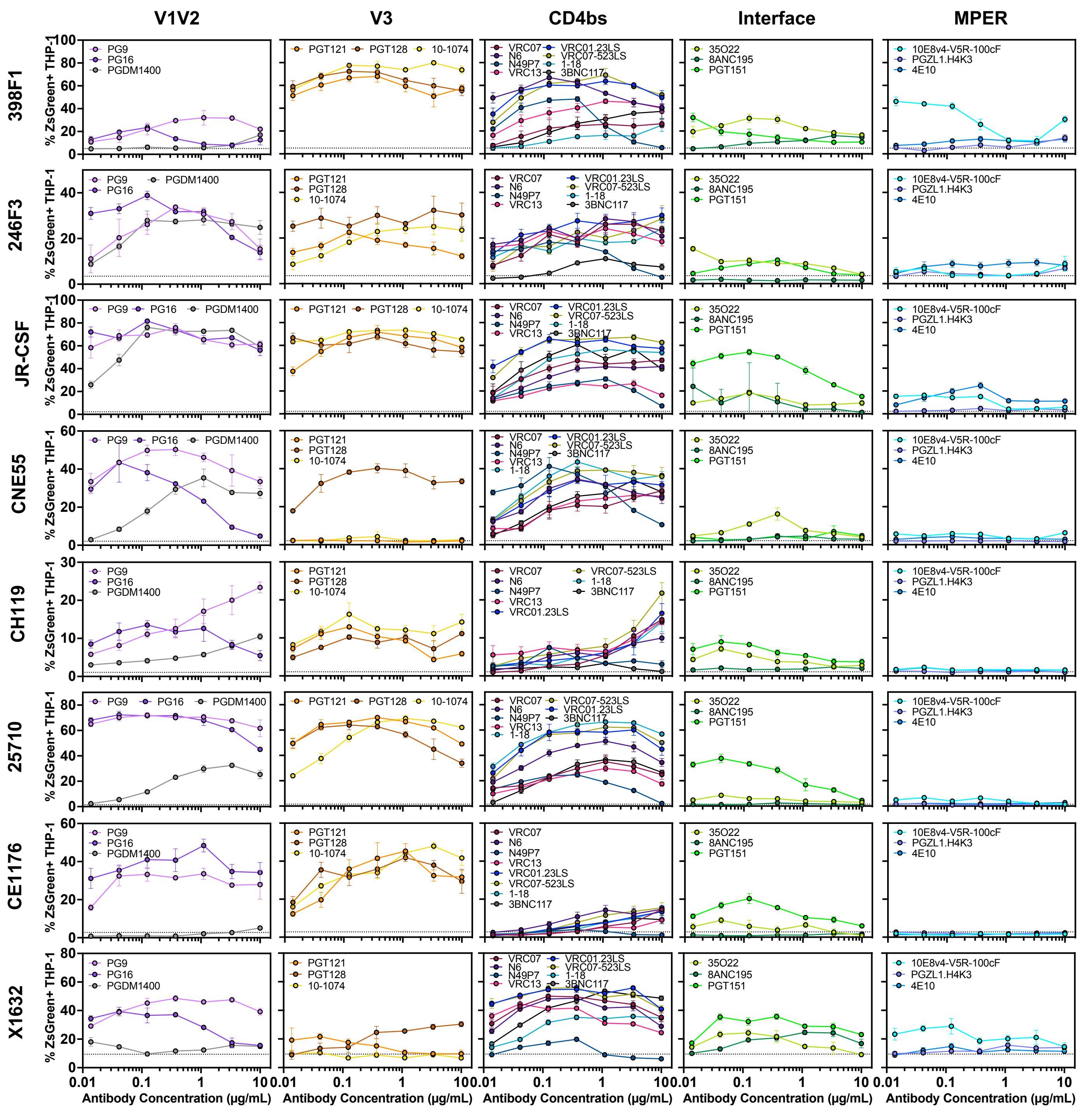


**Figure S10. ADCP of HIV-1 bNAbs to Env-expressing CEM.NKr cells.**
ADCP of twenty HIV bNAbs against CEM.NKr cells expressing each of the diverse HIV-1 Envs, with individual bNAbs indicated in the legends. ADCP activity was performed as mentioned in Fig. S5.

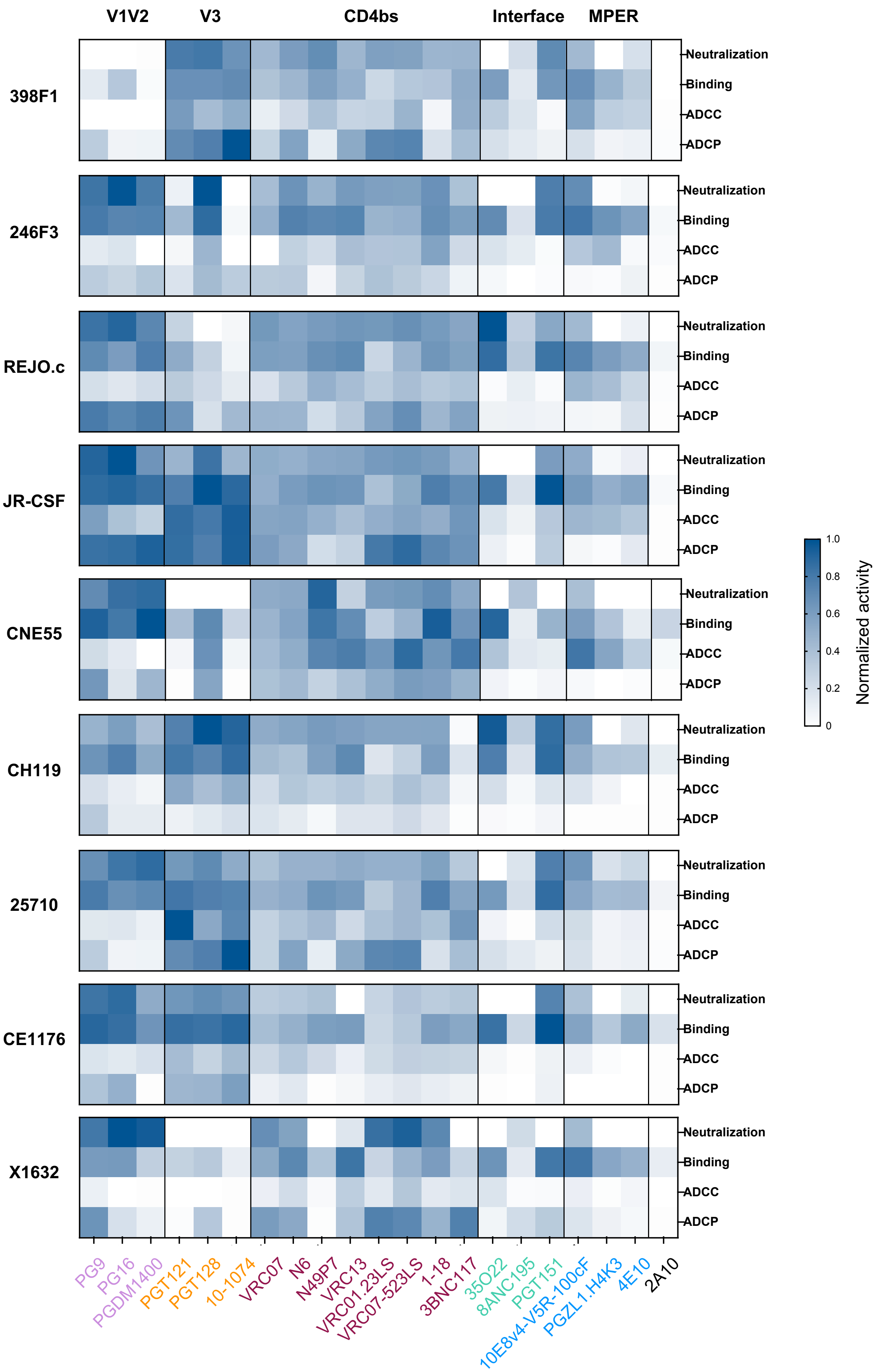
V1V2
V3
CD4bs
Interface
MPER
398F1
246F3
REJO.c
JR-CSF
CNE55
CH119
25710
CE1176
X1632
Neutralization
Binding
ADCC
ADCP
Normalized activity
1.0
0.8
0.6
0.4
0.2
0
PG9
PG16
PGDM1400
PGT121
PGT128
10-1074
VRC07
N6
N49P7
VRC13
VRC01.23LS
VRC07-523LS
1-18
3BNC117
35O22
8ANC195
PGT151
10E8v4-V5R-100cF
PGZL1.H4K3
4E10
2A10

**Figure S11. Functional profiles of bNAbs across diverse Envs.**

Heatmap illustrating the relationship between bNAb-mediated neutralization, Env binding, ADCC, and ADCP for each individual HIV-1 Env. For each Env, bNAbs were assessed for neutralization potency ($–\log_{10}[IC_{50}]$), Env binding (log(AUC)), ADCC (AUC), and ADCP (AUC), and values were independently normalized from 0 to 1 within each function across the HIV-1 panel.

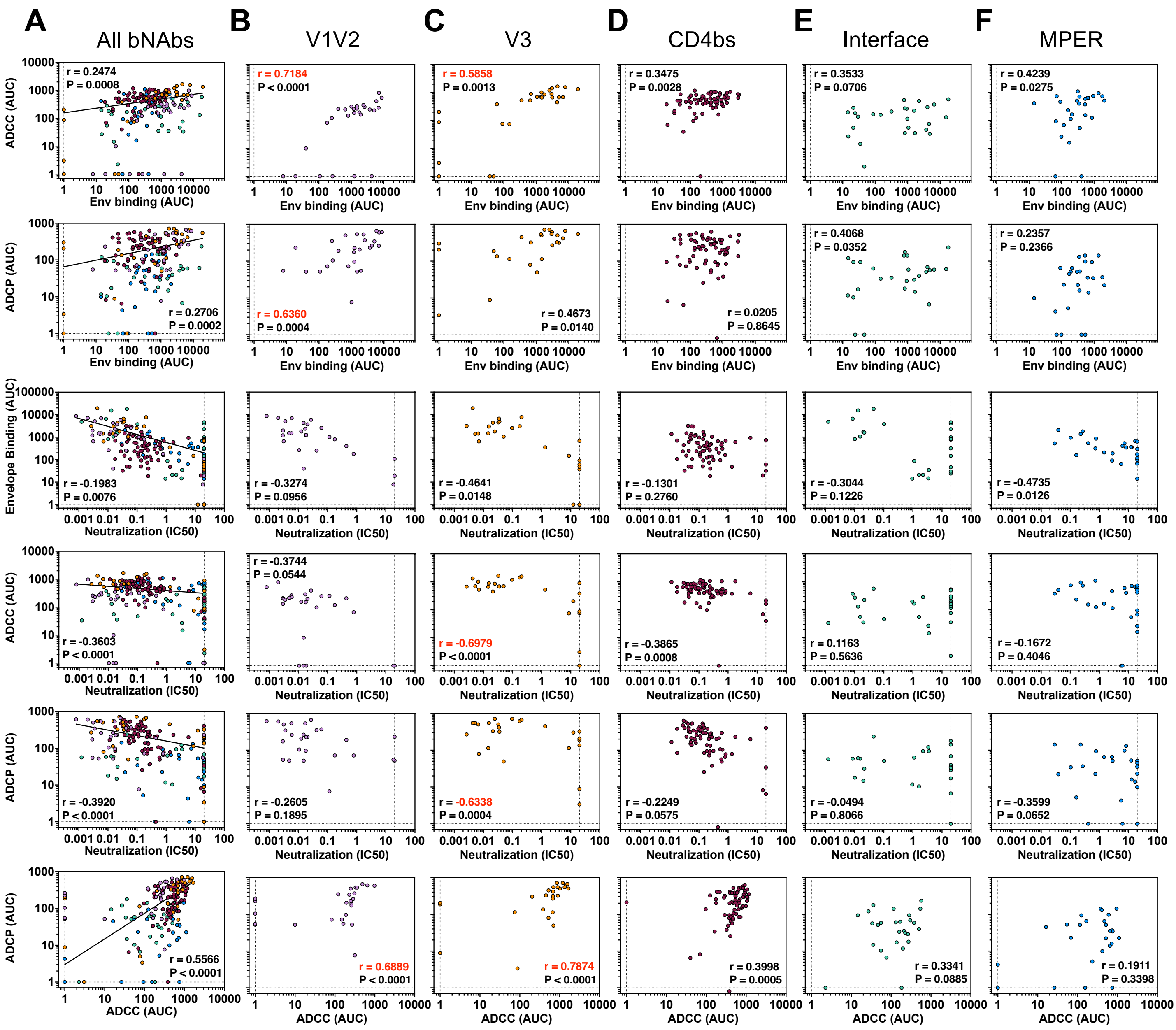


**Figure S12. Distinct correlation patterns of bNAb functions across HIV-1 Env epitopes.**

(**A**) Spearman correlation analyses across all bNAbs evaluating relationships between Env binding, ADCC, ADCP, and neutralization. Each dot represents a single bNAb-strain pair, color-coded by epitope specificity: V1/V2 apex (purple), V3 glycan (orange), CD4 binding site (maroon), gp120/gp41 interface (teal), and MPER (blue). (**B-F**) Correlation analyses performed separately within each epitope class to assess functional relationships among Env binding, ADCC, ADCP, and neutralization.

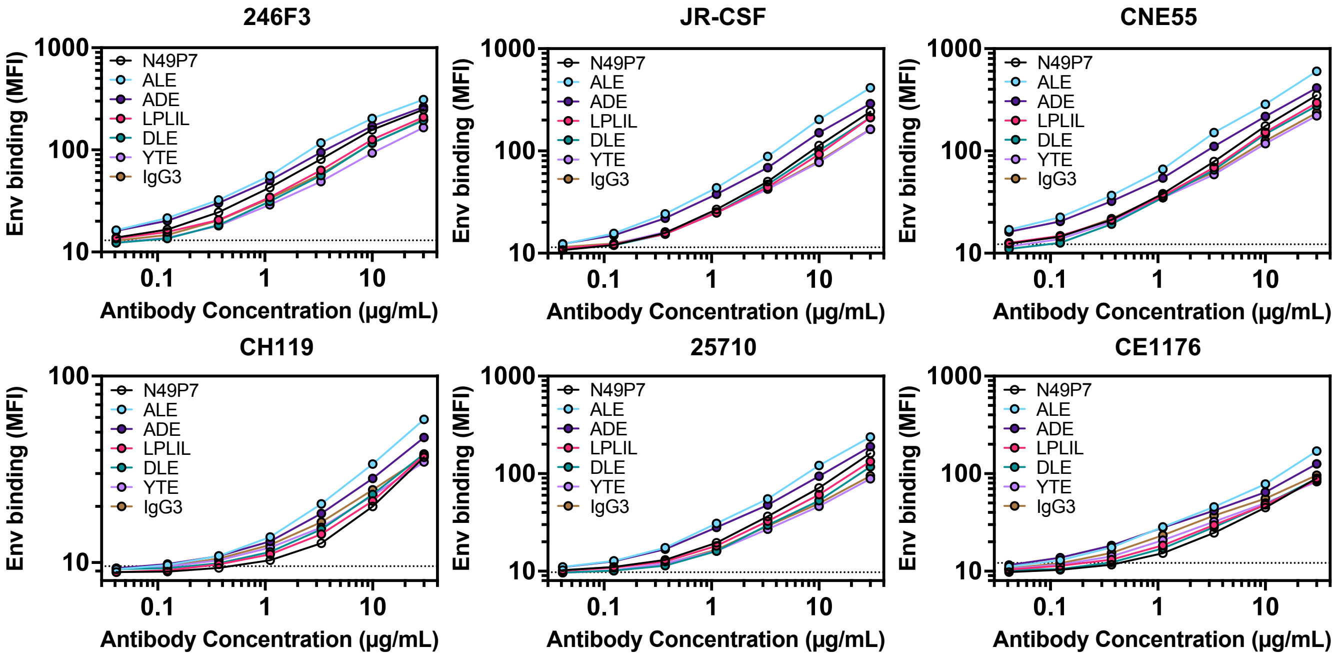


**Figure S13. Fc engineering does not alter N49P7 binding to CEM.NKr cells expressing diverse Envs.**

Binding of AF647-conjugated Fc-engineered N49P7 variants to CEM.NKr cells expressing diverse HIV-1 Envs. Binding is presented as MFI at each bNAb concentration and was performed in triplicate. Dashed lines indicate the baseline activity of the IgG1 control antibody 2A10.

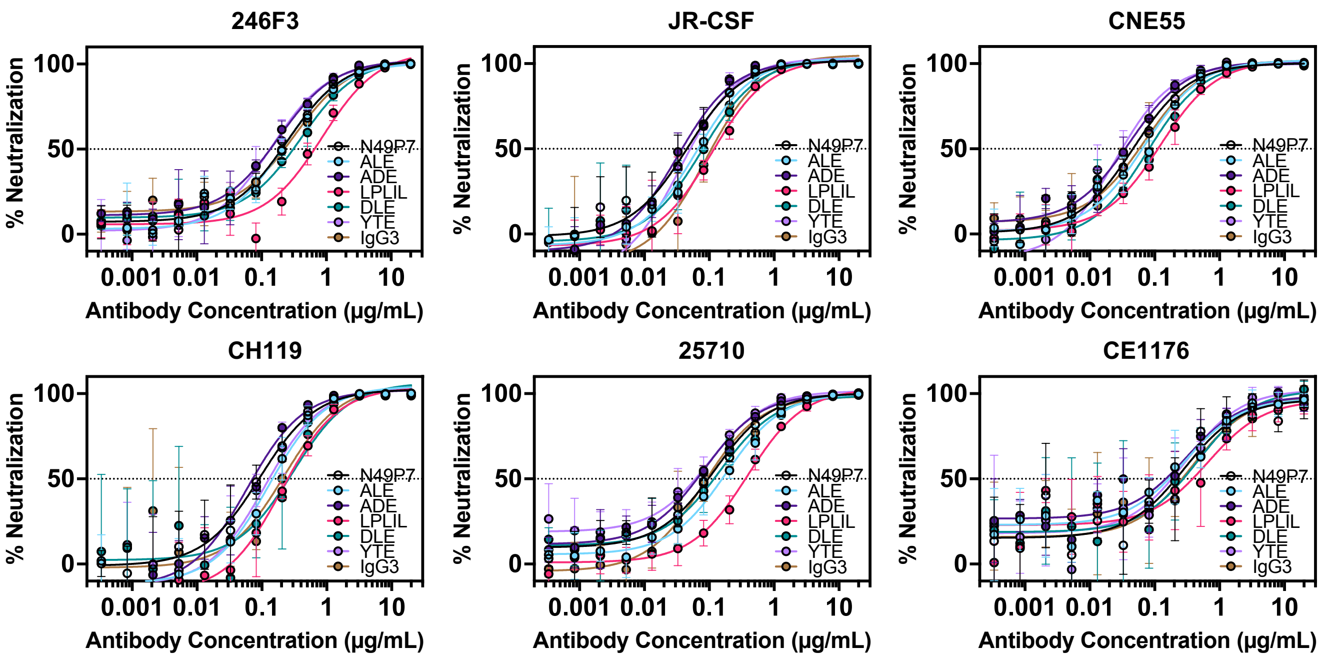


**Figure S14. Fc engineering does not change neutralization activity of N49P7.** Neutralization activity of Fc-engineered N49P7 variants was assessed against diverse HIV-1 strains. Data is shown as percent neutralization across a range of antibody concentrations, with individual bNAbs indicated in the legends.

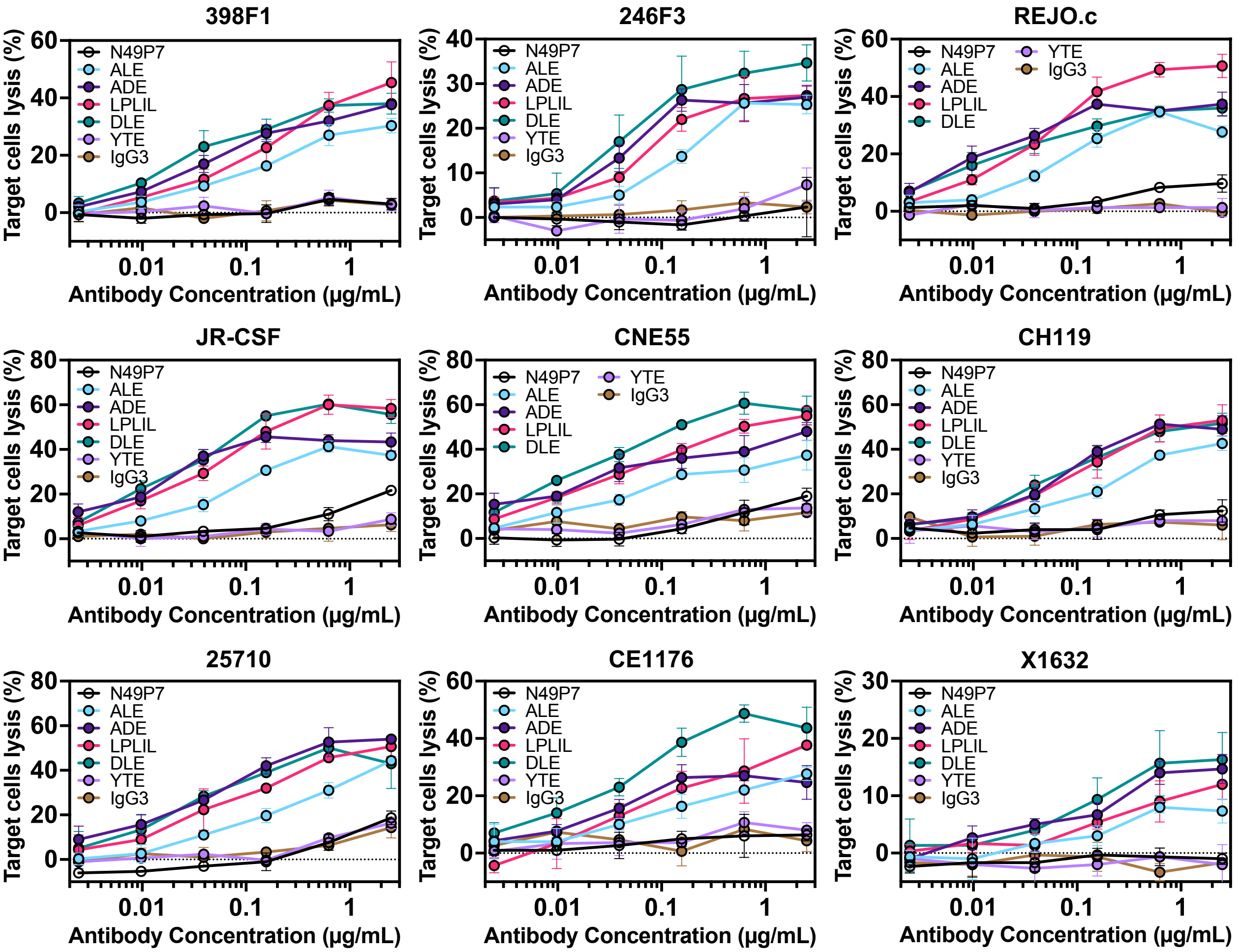


**Figure S15. ADCC activity of N49P7 Fc variants against diverse HIV-1 Envs.**
ADCC activity of N49P7 and its Fc-engineered variants was assessed against diverse Envs. ADCC was quantified as mentioned in Fig. S4. Dashed lines indicate the baseline activity of the IgG1 control antibody 2A10.

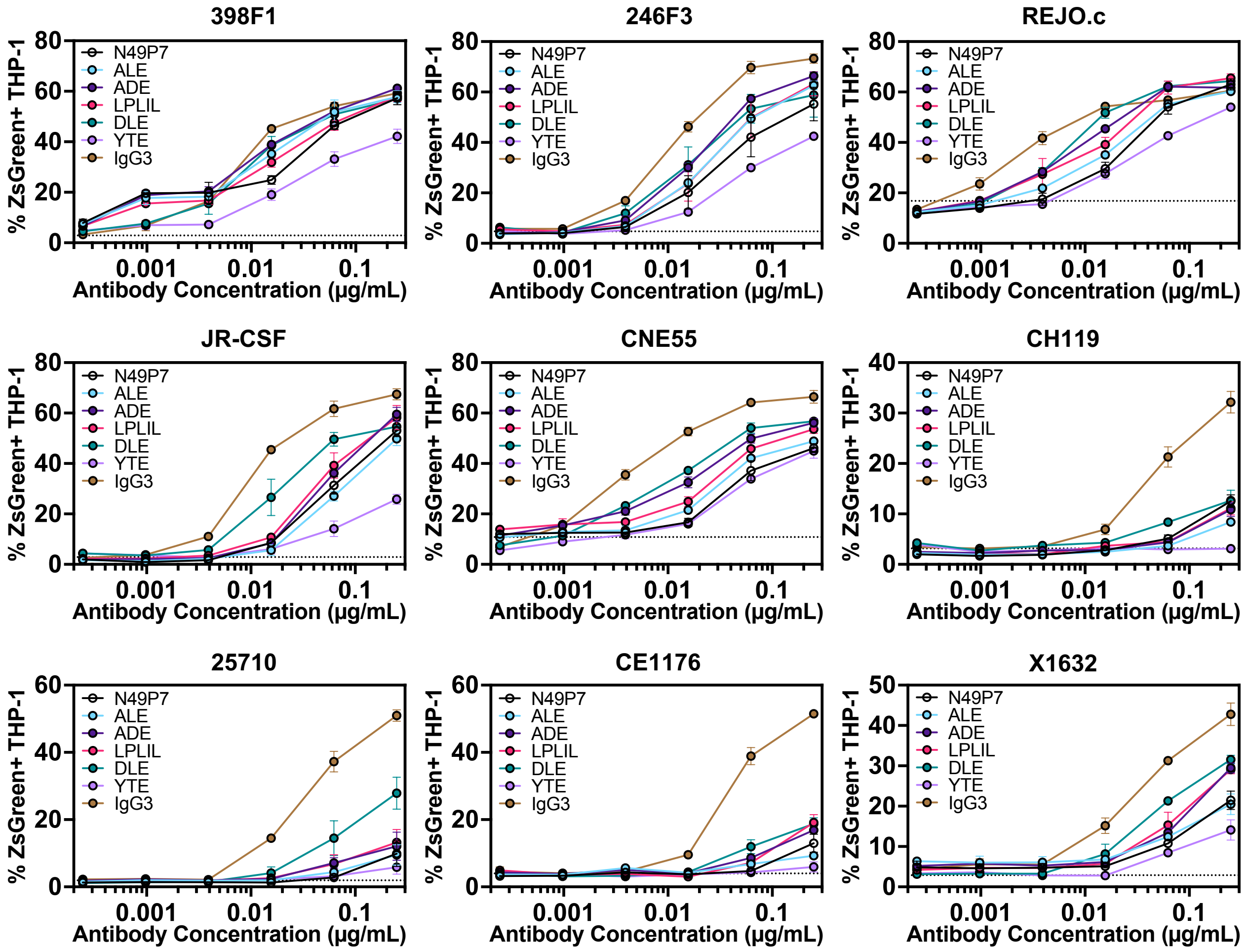


**Figure S16. ADCP activity of N49P7 Fc variants against diverse HIV-1 Envs.**
ADCP activity of N49P7 or its Fc-engineered variants was assessed against diverse Envs. ADCP was quantified as mentioned in Fig. S5. Dashed lines indicate the baseline activity of the IgG1 control antibody 2A10.

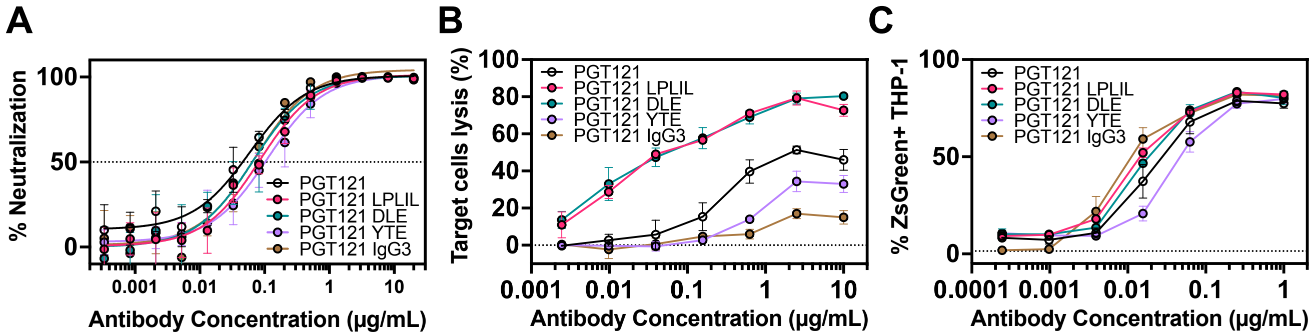


**Figure S17. Fc engineering enhances effector functions of the V3 glycan-targeting bNAb PGT121.**

Functional characterization of wild-type PGT121, Fc-engineered variants, and an IgG3-switched version against $HIV_{JR\text{-}CSF}$. (**A**) Neutralization potency ($IC_{50}$), (**B**) ADCC assay, and (**C**) ADCP assay. Dashed lines in Fig. A indicate 50% neutralization, while in Fig. B and C indicate the baseline activity of the IgG1 control antibody 2A10.
